# Engineering Mechanostable Anticalin Scaffolds to Enhance Particle Adhesion and Targeting of CTLA-4 under Shear Stress

**DOI:** 10.1101/2024.08.16.608260

**Authors:** Yang Sun, Jiajun Li, Rosario Vanella, Haipei Liu, Zhaowei Liu, Michael A. Nash

**Author notes:** These authors contributed equally to this work (Yang Sun, Jiajun Li).

## Abstract

Achieving high binding strength and efficient delivery of molecular cargo to cells expressing biomarker targets is a significant challenge in drug delivery. Here, we investigated how altering the surface immobilization residue (i.e. anchor point) within a non-antibody binding scaffold called Anticalin can enhance particle adhesion to the immune checkpoint protein CTLA-4 on mammalian cells under shear stress. By introducing bio-orthogonal clickable amino acids into Anticalin at various positions and applying tension to the protein complex using single-molecule AFM force spectroscopy and bead-based adhesion assays, we elucidate the relationship between anchor point position and mechanostability of the Anticalin:(CTLA-4) complex. Multi-regression analysis of the physicochemical properties of the anchor points revealed that the distance from the anchor point on Anticalin to CTLA-4’s center of mass was a major determinant of binding strength under shear flow. These results demonstrate how anchor point engineering can enhance particle adhesion and cellular delivery to CTLA-4 targets and provides a heuristic for choosing surface immobilization points of targeting proteins such that they withstand high mechanical forces.

## Introduction

Mechanical anisotropy refers to the difference in mechanical response of a molecule when force is applied from different directions. For instance, double-stranded DNA/RNA helices and hairpins will mechanically unfold at low forces (∼10-20 pN) when unzipping the strands and breaking the H-bonds in series, or exhibit high breakage forces (>50 pN) when loaded in a parallel shear geometry^1, 2, 3^. Similarly, the cooperativity and coordination angles of chelated metal ions impart high mechanical stability (∼220pN) when tension is applied along particular a pulling direction to 3WJ-pRNA^4, 5^. Folded protein domains also respond to mechanical force differently depending on the pulling geometry, as was shown for several globular domains (e.g., GFP^6^, GB1^7, 8^, ubiquitin^9^, and E2lip3^10^). Similarly, the strength of protein-protein complexes subjected to mechanical force depends on whether the domains are anchored from the N- or C-terminus, as was shown in streptavidin-biotin and cohesin-dockerin systems^11, 12, 13^. Differences in shearing versus unzipping geometries of protein coiled-coils have also been reported^14^. Despite these anecdotal studies on mechanical anisotropy of protein domains, we currently lack design rules or heuristics for achieving mechanically stable protein-protein interactions, and it remains unclear how pulling geometry modulates the binding strength of protein-protein complexes under conditions of flow and force.

The influence of mechanical forces on protein-protein interactions extends beyond the molecular level, reaching into the realm of nano-, biological and soft matter particles (e.g., polymeric/hydrogel particles, liposomes, viral particles, etc…) which are being investigated as drug delivery vehicles and molecular imaging probes. These particles may encounter complex fluidic forces in the bloodstream which can shear them off of cell surfaces. Indeed, the adhesion of targeted nanoparticles and microbeads and their delivery efficiency to cells is known to severely diminish in the presence of shear flow.^15, 16, 17^

To tackle this challenge, here we have developed an approach for studying and optimizing anchor points within a binding scaffold in order to enhance the mechanical stability of the complex with its target ligand, and thereby improve the cellular adhesion strength of receptor-targeted particles. In this context, “anchor point” refers to the precise amino acid residue number within the primary sequence used to immobilize the targeting protein domain onto the particle surface. Specifically, we investigated a non-antibody binding scaffold called Anticalin that forms a complex with the immune checkpoint protein cytotoxic T-lymphocyte-associated protein 4 (CTLA-4, also known as CD152) ^18,19, 20^. The Anticalin we investigated was previously engineered from human Lcn2 to specifically bind CTLA-4^21^. With its stimulatory activity on T cells, this Anticalin and similar molecules have potential applications in targeted immunotherapy.

We used AFM-based single-molecule force spectroscopy and quantitative bead-based adhesion assays to study force-induced dissociation of the engineered Anticalin from its target (CTLA-4) under various pulling geometries. By pulling Anticalin from 17 different anchor points, we mechanically disrupted the interaction with CTLA-4 and studied how the choice of anchor point altered mechanostability of the protein-protein complex. We selected anchor points embedded within different secondary structures and possessing different biophysical properties, and correlated these properties with the mechanostability of the complex. This analysis provides for the first time guidelines for anchor point selection with potential to be generalized to other binding scaffolds. We further validated the concept at the cellular scale by quantifying adhesion of Anticalin-modified microparticles to CTLA-4^+^ cells under the influence of shear flow, and found that functionalizing the particles through Anticalin’s mechanostable anchor point was sufficient to improve particle adhesion. Our results provide insight into the dissociation pathways of protein complexes under different loading geometries, demonstrate a mechanical analog to traditional affinity maturation, and provide design parameters for enhancing binding strength through rationale choice of anchor points within scaffolds.

## Results and Discussion

### Rationale design of Anticalin pulling points based on structure

We first sought to design positions within the Anticalin sequence where we could introduce bio-orthogonal clickable azides to apply mechanical force to the Anticalin:(CTLA-4) complex. Lipocalins share a highly conserved 8-stranded anti-parallel β-barrel with α-helices along the sides, a ligand-binding open end and a more compact closed end^22^. The structure of the Anticalin complexed with the extracellular domain of CTLA-4 is shown in **Fig. 1** (PDB 3BX7)^21^. In protein secondary structures, beta strands primarily act as force-bearing rigid units, while alpha helices are typically more flexible^23, 24^. Based on Anticalin’s structural features, we rationally chose a series of 17 anchor points spanning the entire protein sequence length (**Fig. 1a, b**, and **Table S1**). Using single-molecule AFM, we first studied the mechanical stability of the Anticalin:(CTLA-4) complex by pulling Anticalin from these different anchor points. Each anchor point on the Anticalin molecule corresponds to a different Anticalin variant where a bio-orthogonal azide was introduced to achieve the precisely defined pulling geometry in the AFM measurements, as well as the bead-based adhesion measurements (**Fig. 1c**).

**Fig. 1:**
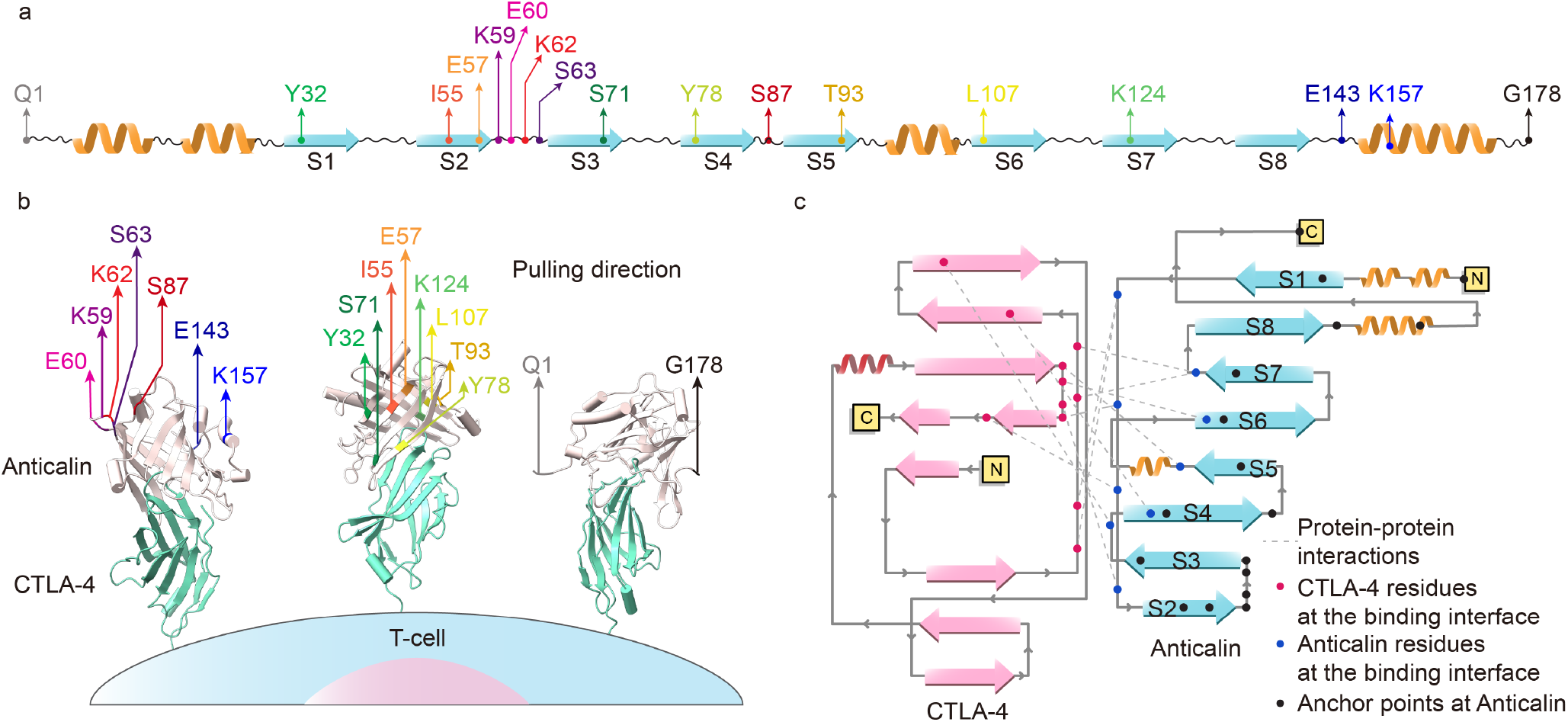
Anchor points selection in Anticalin. **a**, Schematics of Anticalin secondary structure and the distribution of chosen anchor points across secondary structures (i.e. linkers, α helixes, and β-strands). **b**, Schematic of Anticalin tertiary structure and anchor point distribution. Structure of CTLA-4 in complex with Anticalin (PDB 3BX7). The pulling directions on the Anticalin are indicated by colored arrows, which originate from the anchor points. The anchor point on CTLA-4 is fixed at the C-terminus, replicating the natural tethering geometry found on the T-cell surface. **c**, Schematic of Anticalin:(CTLA-4) complex secondary structure. Anticalin features a central β-barrel composed of eight antiparallel β-strands (S1–S8), which are interconnected by short, flexible linkers. The anchor points, indicated by black dots, are rationally designed at the closed end of the β-barrel to minimize interference with the binding interface. Red dots represent CTLA-4 residues at the binding interface, blue dots represent Anticalin residues at the binding interface, and grey dashed lines indicate protein-protein interactions (salt bridges and hydrogen bonds) at the binding interface.

### Single-molecule analysis of anchor point-dependent mechanostability

To connect Anticalin to the AFM tip through a given internal anchor point, we combined non-canonical amino acid (ncAA) incorporation, clickable peptide handles, site-specific surface immobilization, and an AFM-SMFS measurement setup with freely diffusing receptor molecules that was previously reported by us and others^11, 12, 25^ (**Fig. 2a**). Azide groups were introduced at different positions in Anticalin through amber codon suppression to replace the selected anchor point residue with azido-phenylalanine. Subsequently, synthetic peptides consisting of the N-terminal fibrinogen β (Fgβ) sequence followed by a Strep tag and a C-terminal dibenzocyclooctyne (DBCO) group were conjugated to the azide group on the respective Anticalin variant using copper-free click chemistry. Each Anticalin variant containing Fgβ attached through a different anchor point was purified using size exclusion and Strep-trap columns to remove excess peptides and unreacted Anticalin. SDS-PAGE and mass spectrometry analysis confirmed the successful conjugation of Fgβ, increasing the molecular weight of the respective Anticalin by 3 kDa (**Fig. S1**). Each Anticalin with Fgβ clicked onto the given anchor point residue was then measured in separate AFM experiments.

**Fig. 2:**
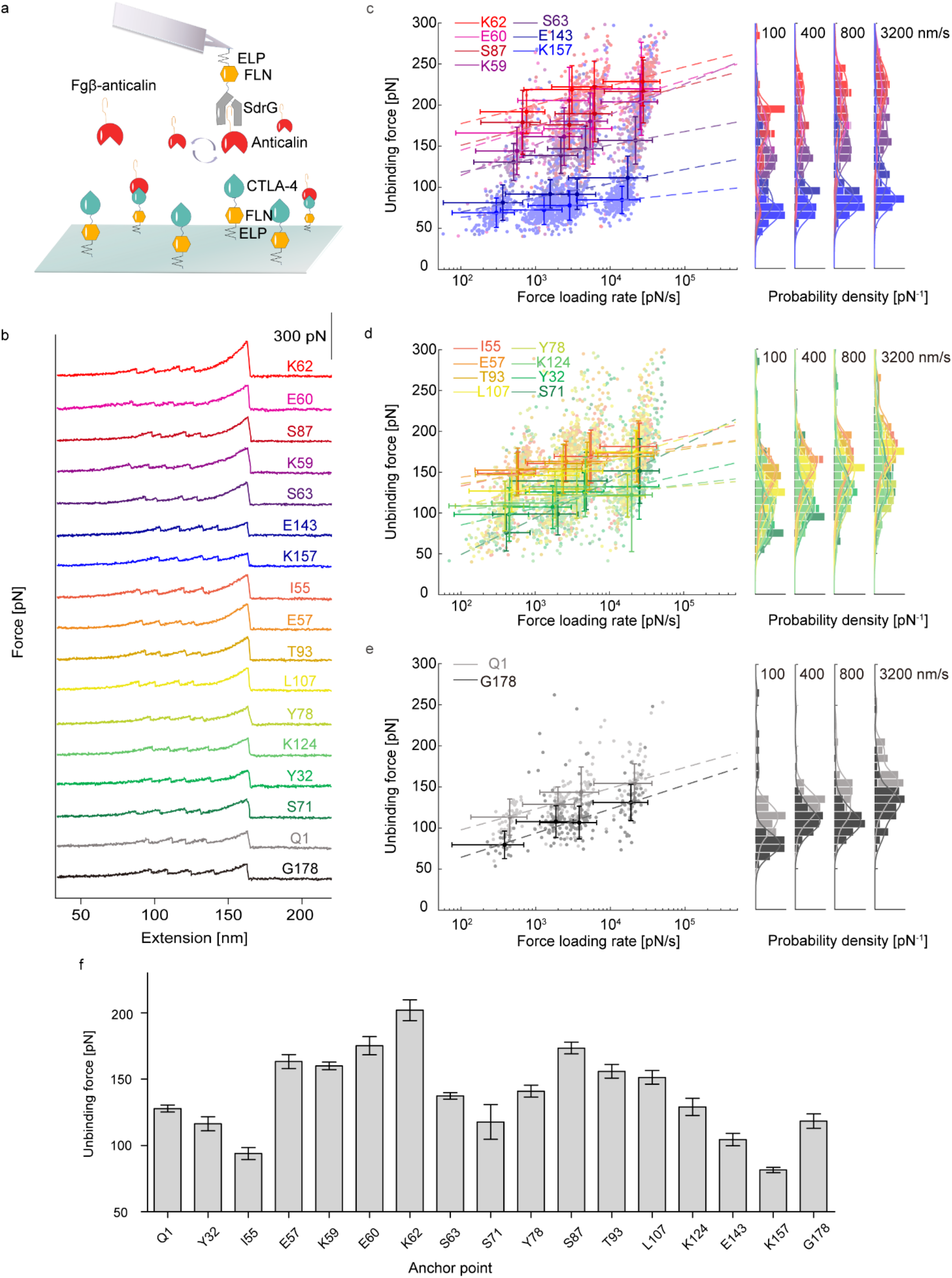
Single-molecule analysis of anchor point-dependent mechanostability. **a**, Schematics of AFM experiments. **b**, AFM curves of rupture events between Anticalin and CTLA-4 under different unbinding geometries. **c-e**, Most probable rupture forces measured at different pulling speeds were plotted against the logarithm of average loading rate and fit linearly to extract the zero-force off rate *k*_*0*_ (BE) and distance to the energy barrier Δx (BE). Error bars represent the standard deviation of rupture forces and loading rates. **c**, anchor residues at loops and α helix, **d**, β sheets, **e**, N/C terminal. **f**, Most probable rupture forces between Anticalin and CTLA-4 when pulled at 400 nm/s were plotted against the anchor residue number on Anticalin. Error bars represent the standard error of rupture forces measured at 400 nm/s.

For the AFM setup, CTLA-4 and SdrG were first immobilized onto the cantilever and substrate through site-specific attachment *via* vbbR tags, respectively. The Anticalin with Fgβ peptide conjugated at the selected anchor point was added to the measurement buffer to achieve a final concentration of 1 μM. When the cantilever approached the substrate, the Fgβ at one end of the freely diffusing Fgβ-Anticalin construct could bind to SdrG attached to the cantilever, while the other end (i.e. the open end of Anticalin) could bind to CTLA-4 immobilized on the substrate, forming a ternary complex. After lifting the cantilever, dissociation events between Anticalin and CTLA-4 were observed, with unfolding events of two FLN structural domains serving as fingerprint patterns for identification of valid single-molecule traces (**Fig. 2b**). In a typical overnight AFM measurement, about ∼10,000 force-extension curves were recorded and converted to contour length space using the freely rotating chain (FRC) elasticity model. Curves were filtered to identify traces with the two-step unfolding pattern of two FLN fingerprint domains and 2x 32 nm contour length increments. The SdrG:Fgβ complex could withstand forces up to 2 nN. When the cantilever was retracted, the significantly weaker Anticalin:(CTLA-4) complex dissociated first, leaving the Fgβ-Anticalin on the cantilever. The moderate equilibrium affinity between SdrG and Fgβ (K_D_ ∼ 400 nM) allowed for the rapid exchange of Anticalin molecules on the cantilever. In this setup, the cantilever tip and surface molecules are always newly probed, thus the re-foldability of the cantilever molecules does not play a role. The forces involved in the dissociation events between Anticalin and CTLA-4 were then analyzed to quantify the mechanical stability between CTLA-4 and Anticalin when pulled from different loading points.

Using this setup, we measured the rupture forces of the Anticalin:(CTLA-4) complexes at four pulling speeds ranging from 100 to 3200 nm/s, and plotted the rupture force distributions as histograms (**Fig. 2c-e**). We fit the histograms to extract the most probable rupture forces and plotted them against the anchor point residue number on Anticalin, revealing intriguing trends on how the mechanostability of the Anticalin:(CTLA-4) complex correlates with structural features and physicochemical properties of the anchor points. We initially categorized different anchor points based on their secondary structure elements into four groups: Loops, α-helices, β-strands, and N/C-termini. For a clear and direct comparison, anchor points within loops and α-helices, which are less resistant to force, were visualized in **Fig. 2c**; those within the force-bearing β-strand structures are shown in **Fig. 2d**; and those at the N/C termini are in **Fig. 2e**. Under different unbinding geometries, the dissociation of Anticalin:(CTLA-4) complexes exhibited significant mechanical anisotropy. When stretched from the K157 anchor point at a pulling speed of 400 nm/s within alpha helix #3, the complex showed the weakest mechanical stability (81.5 ± 1.9 pN, Mean ± SE) of all pulling geometries. Meanwhile, stretching from N- or C-terminal (Q1 or G178) anchor points generated medium to low stability (127.8 ± 2.6 pN, and 118.4 ± 5.4 pN). Notably, stretching from the K62 anchor point within a central loop showed the highest rupture force across all pulling geometries (201.9 ± 7.8 pN).

As shown in **Fig. 2f**, anchor points in beta strand 1: residue E57 (163.2 ± 5.3 pN) and those located in the loop between beta strands 2 and 3: residues K59 (160.0 ± 2.9 pN), E60 (175.2 ± 6.9 pN), and K62 (201.9 ± 7.8 pN) generated higher complex stability. Other stable points were found in the loop bridging beta strands 4/5: S87 (173.4 ± 4.4 pN), and within beta strands 5 and 6: residues T93 (155.9 ± 5.2 pN) and L107 (151.4 ± 5.1 pN). In contrast, anchor points in beta strands 1 and 3: residues Y32 (116.4 ± 5.4 pN), I55 (94.0 ± 4.5 pN) and S71 (117.8 ± 13.1 pN), and in the C-terminal alpha helix: residue K157, as well as N- and C-terminal anchor points generated low mechanostability. T-tests were performed to analyze significant differences in mechanostability between any 2 anchor points (**Table S2)**.

### Anchor point dependent energy landscapes

Next, we used the Bell-Evans (BE) and Dudko-Hummer-Szabo (DHS) models to estimate the energy landscape parameters for each pulling geometry^26, 27, 28, 29, 30^. As shown in **Fig. 2c-e**, the most probable rupture forces were linearly fit against the logarithm of the average loading rate at a given pulling speed to extract the zero-force off-rate (*k*_*0*_) and the distance to the energy barrier (Δx) using the BE model. Subsequently, using the DHS model we transformed the rupture force histograms into force-dependent lifetimes, applying previously reported fitting methods to obtain τ_0_, Δx, and the height of the energy barrier (ΔG)^26^. **Table S4** lists the parameters extracted from both models. According to **Fig. 2, Fig. S3**, and **Table S4**, the energy landscapes measured from the different pulling geometries possess energy barriers of varying shapes and heights. Notably, mechanically stable pulling geometries often have relatively higher energy barriers ΔG, or shorter distance to the transition state Δx, contributing to greater resistance to external forces. In contrast, geometries with lower mechanical stability tended to have lower energy barriers or longer Δx. Thus, the mechanical stability of the complex is determined by the interplay between the height and shape of the energy landscape, which depends on the loading geometry. Moreover, **Fig. S3** shows that depending on the anchor point, the bound lifetime τ_off_ at a given force varied by 3-4 orders of magnitude. For example, the log (τ_off_) value of Anticalin K62 is -0.9, in contrast to the log (τ_off_) value of Anticalin K157, which is -4.5 (**Fig. S3**, 100 pN).

### Correlating mechanostability with biophysical properties of anchor points

To understand these effects more deeply, we parameterized structural and physicochemical properties of the anchor points and sought to draw correlations with the rupture force data. The parameters included in this analysis were hydropathy, electronic charge index, solvent accessible surface area (SASA), and distance from the Anticalin anchor point to the CTLA-4 center of mass^31, 32^. Initially, we attempted to fit the mechanical stability of all anchor points as a linear combination of these parameters, generating a line fit with R^2^ value of 0.470. The importance of each parameter was reflected by its feature score. The higher the feature score, the more relevant this parameter was to the complex mechanical stability. As shown in **Fig. S4a**, the feature score of distance to the center of mass of CTLA-4 is at least 4.8 times higher than other parameters, making it the most decisive factor. To further explore the correlation between each parameter and the mechanical stability of the complex, we performed linear regression analysis on each parameter against mechanical stability, respectively. As shown in **Fig. S4b-e**, the distance to the center of mass of CTLA-4 has an R^2^ value of 0.408, much higher than that of 0.003, 0.013, and 0.125 for hydropathy, electronic charge index, and SASA, respectively.

Next, considering different secondary structures of proteins have distinct responses to mechanical forces, we hypothesized that anchor points located in different secondary structure regions might behave differently. We separated the anchor points into groups according to their secondary structural labels (e.g., being located in loops, alpha helices, or beta strands). Then we repeated the above steps for multiparametric linear fitting of all parameters together. This achieved a higher fitting accuracy with an R^2^ value of 0.777 for the group with loops and alpha helices, and an R^2^ value of 0.760 for the group with beta strands. The factor most significantly affecting mechanical stability was still the distance to the CTLA-4 center of mass, as demonstrated by its highest feature score (**Fig. S5a** and **S6a**). Further validating the correlation by fitting the unbinding force separately with each physicochemical parameter confirmed our findings that the mechanical stability of the complex is related to the distance between the anchor point in Anticalin from the center of mass of CTLA-4. As can be shown in **Fig. S5b-e** and **S6b-e**, the distance to the center of mass of CTLA-4 still has the highest R^2^ value among all parameters when fitting with the unbinding force. The mechanical stability increases as the distance to CTLA-4 increases, and is unrelated to hydropathy, electronic charge index, or solvent accessible surface area of the anchor point. Furthermore, when linearly fitting the distance to the center of mass of CTLA-4 and the mechanical stability of the complex, anchor points located in loops and alpha helices exhibited higher R^2^ value than those in beta strands, highlighting the impact of the secondary structure region where the anchor point is located on the overall stability of the complex. To our knowledge, the finding that anchor points located far away from the center of mass of the bound ligand can generate high levels of mechanical resistance in molecular complexes represents a new and previously unreported design heuristic for engineering molecular complexes with high mechanical stability. We further compared the individually fitted energy landscape parameters to the distance between the anchor point and CTLA-4 center of mass, but found no significant correlations (**Fig. S7**).

### Analysis of equilibrium binding affinity by thermophoresis

We next tested the binding affinity at equilibrium (i.e. in the absence of force). We measured the equilibrium dissociation constants (K_D_) between CTLA-4 and four of our Anticalin mutants using Microscale Thermophoresis (MST). As illustrated in **Fig. S2** and **Table S3**, despite significantly different mechanical responses of the Anticalins with an azide installed at positions Q1, K62, K157, or G178, they all exhibit very similar affinities to CTLA-4 at equilibrium, showing no significant differences. This is not surprising because the introduced pulling handles were located on the opposite side of Anticalin as the binding epitope for CTLA-4 and presumably did not significantly alter the binding interface.

### Mechanostable anchor points enhance bead adhesion to CTLA-4^+^ cells

Particle adhesion strength to cells is governed by the resistance of the receptors to mechanical forces transmitted from the shear fluid to the beads. We hypothesized that immobilizing Anticalin through stable anchor points could enhance bead adhesion. To demonstrate this concept, we worked with PolyAn beads (∼20 µm diameter) consisting of poly(methyl methacrylate) (PMMA) cores coated with poly(acrylic acid) (PAA) shells modified with DBCO groups on the surface^33, 34^. Utilizing the same bioconjugation strategy as in the AFM experiments, we conjugated Anticalins bearing a single non-canonical azide group at the desired anchor point to the DBCO group on the PolyAn beads using copper-free click chemistry. To determine the amount of each azide-bearing Anticalin attached to the polybeads, we performed DBCO labeling and surface passivation (see methods), followed by fluorescent immunostaining of the 6x-Histidine tag present at the terminus of each Anticalin. We then quantified the degree of bead modification using flow cytometry (**Fig. S8**). After antibody labeling, the successfully modified beads generated a green fluorescence signal on the surface, and the intensity of the fluorescence signal was positively correlated with the amount of Anticalin used during the surface modification step (**Fig. S8a**). This analysis allowed us to achieve similar bead modification levels for different Anticalin variants (equal avidity case), despite differences in reactivity among the Anticalin variants with azides in different sequence positions. This was achieved by adjusting the concentration of the respective Anticalin during the surface conjugation reaction to achieve very similar fluorescence intensity values for all bead samples (**Fig. S8b**). In parallel, we prepared beads that were maximally modified with Anticalin, without regard for differences in degree of modification among the different anchor points (maximal avidity case) (**Fig. S8b**). Using these two bead conditions (equal avidity or maximal avidity), we then evaluated the binding efficiency of the beads to CTLA-4^+^ CHO-K1 cells under various forms of fluid shear stress, and measured bead adhesion as a function of anchor point residue.

We chose residues Q1, K62, K157, and G178 as the anchor points in the bead adhesion study. K62 and K157 showed the highest and lowest mechanical stability, respectively, in the AFM experiments, while positions Q1 and G178 represent standard tethering from N- and C-terminal ends respectively. Moreover, our MST experiments demonstrated that Anticalins bearing azides at these positions all had similar affinity to CTLA-4 at equilibrium (**Fig. S2**).

For the bead adhesion study, CTLA-4^+^ CHO-K1 cells were seeded into a 48-well plate. When the cells reached ∼90% confluency, an equal number of polybeads with equal or maximized Anticalin avidity were added to each well. We applied mechanical stimulation using a rotary shaker set at 300, 600, or 900 RPM for 1 hour, followed by washing, replacing with fresh culture medium, and continuing shaking for an additional hour. The adhesion strength of the bead samples was evaluated using fluorescence microscopy and image analysis, by dividing the number of adhered beads after the shaking/washing protocol by the total number of live cells. Our results indicated that in the absence of external forces (i.e. no shaking), the binding efficiency of all bead samples was essentially similar, with no significant differences observed depending on the Anticalin anchor point. Under a small mechanical stimulus of 300 RPM, we also found no significant differences in binding efficiency between force and no force conditions. Conversely, the mechanical stimulus applied at 900 RPM was too strong, resulting in very low binding efficiency for all bead samples to CTLA4^+^ cells.

We then used 600 RPM as an intermediate level of mechanical stimulation for the bead adhesion assays. As shown in **Fig. 3**, when the beads were modified with an equivalent avidity of Anticalin through different anchor points, the K62 anchor point demonstrated the highest binding efficiency to CTLA-4^+^ cells, which was also consistent with the results from single-molecule observations. No significant differences were observed among the other anchor points, which could be due to the mechanical stability of the other anchor points being too weak, or a generally lower resolution/sensitivity of the bead adhesion assays as compared with AFM. When the beads were saturated with Anticalin without regard to differences in degree of bead modification (i.e. maximal avidity), the K62 anchor point again showed the highest binding efficiency to CTLA-4^+^ cells compared to anchor points K157 and G178. However, under this condition no significant difference was observed when comparing K62 to anchor point Q1. We note that the Q1 anchor point under the maximal avidity condition was found to have 1.4-fold more Anticalin on the bead surface (**Fig. S8b**) than K62. This result demonstrates how receptor mechanostability and avidity both play a role in determining bead adhesion strength under flow.

**Fig. 3:**
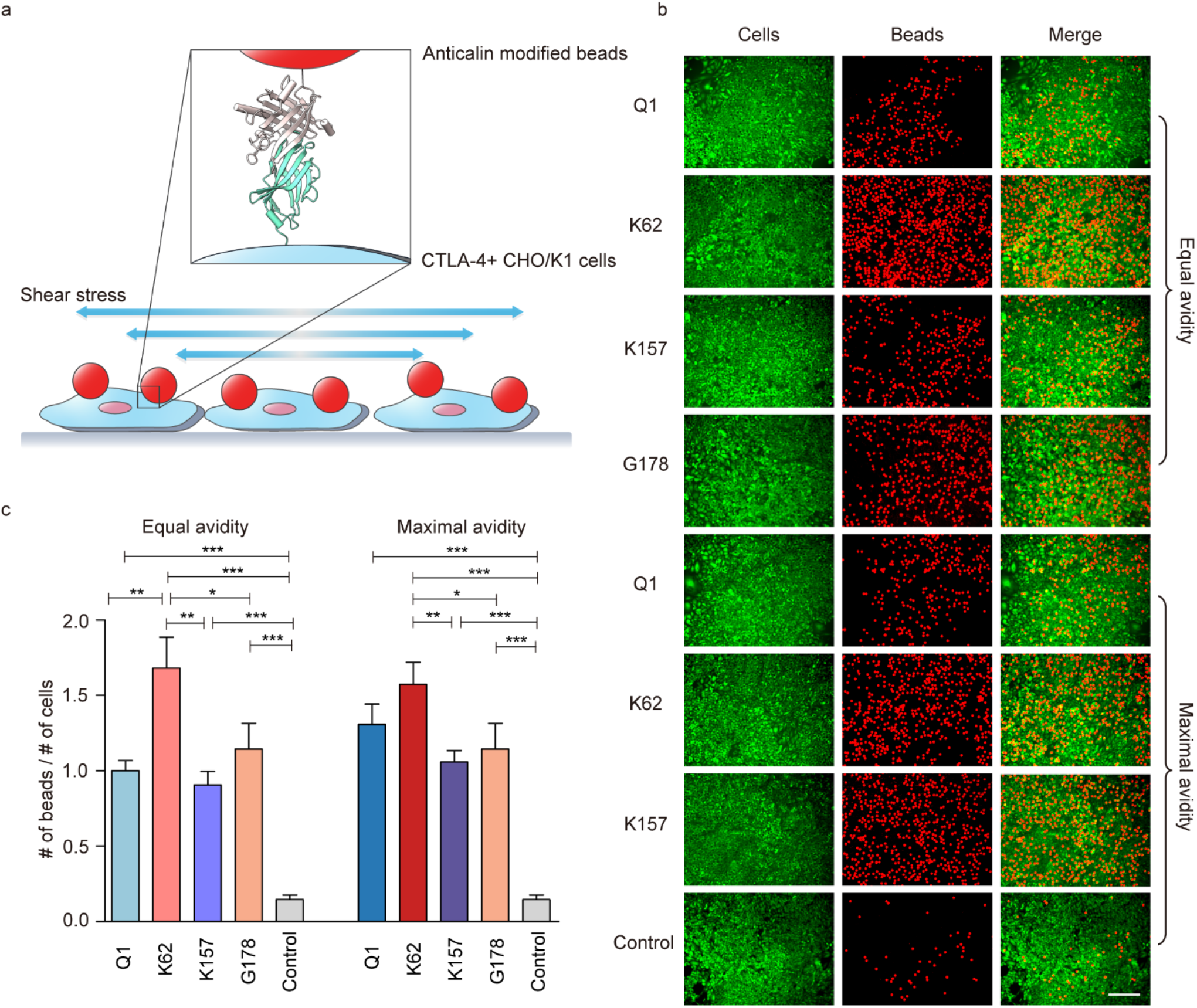
Bead adhesion to CTLA-4+ cells. **a**, Schematics of shaking culture experiments. Anticalin modified beads were allowed to bind to CTLA-4+ CHO/K1 cells under shear stress introduced by rotary shaking during culture. **b**, Fluorescence microscopy images of CTLA-4+ CHO/K1 cells (green) and polybeads (red), scale bar, 200 µm. **c**, Binding affinity between Anticalin with different amber mutation sites and CTLA-4+ cells were characterized by dividing number of polybeads by population of cells, the data was further normalized by Anticalin Q1 at equal avidity group. Statistically significant differences are indicated as *P < 0.05, **P < 0.01, and ***P < 0.001.

### Shear stress assays to quantify bead adhesion to CTLA-4^+^ cells

Next, we developed a quantitative shear stress assay based on a rotating coverglass disk to study anchor point-dependent bead adhesion^35^. CTLA-4^+^ CHO-K1 were cultured on amino-silanized cover glasses in growth medium. Similar to the shaking culture experiments, beads modified with Anticalin through different anchor points (Q1, K62, K157, and G178) were incubated with CTLA-4^+^ cells. After removing the medium containing unbound beads, the substrates were transferred to a spinning disk sample holder/cassette (**Fig. 4a**). Using a spinning protocol, cover glasses were exposed to a gradient of shear stress that grew linearly from the center to the edge. We note that the centrifugal forces in this assay are negligible, and the dominant forces on the cells are due to shear flow parallel to the surface plane and pointing radially outward from the disk center. Under this shear stress, Anticalin-modified beads detached from the surface of the CTLA-4^+^ cells. As the shear stress increased along the radius, more beads detached from the cells, resulting in a lower bead density at the edge of the disk.

**Fig. 4:**
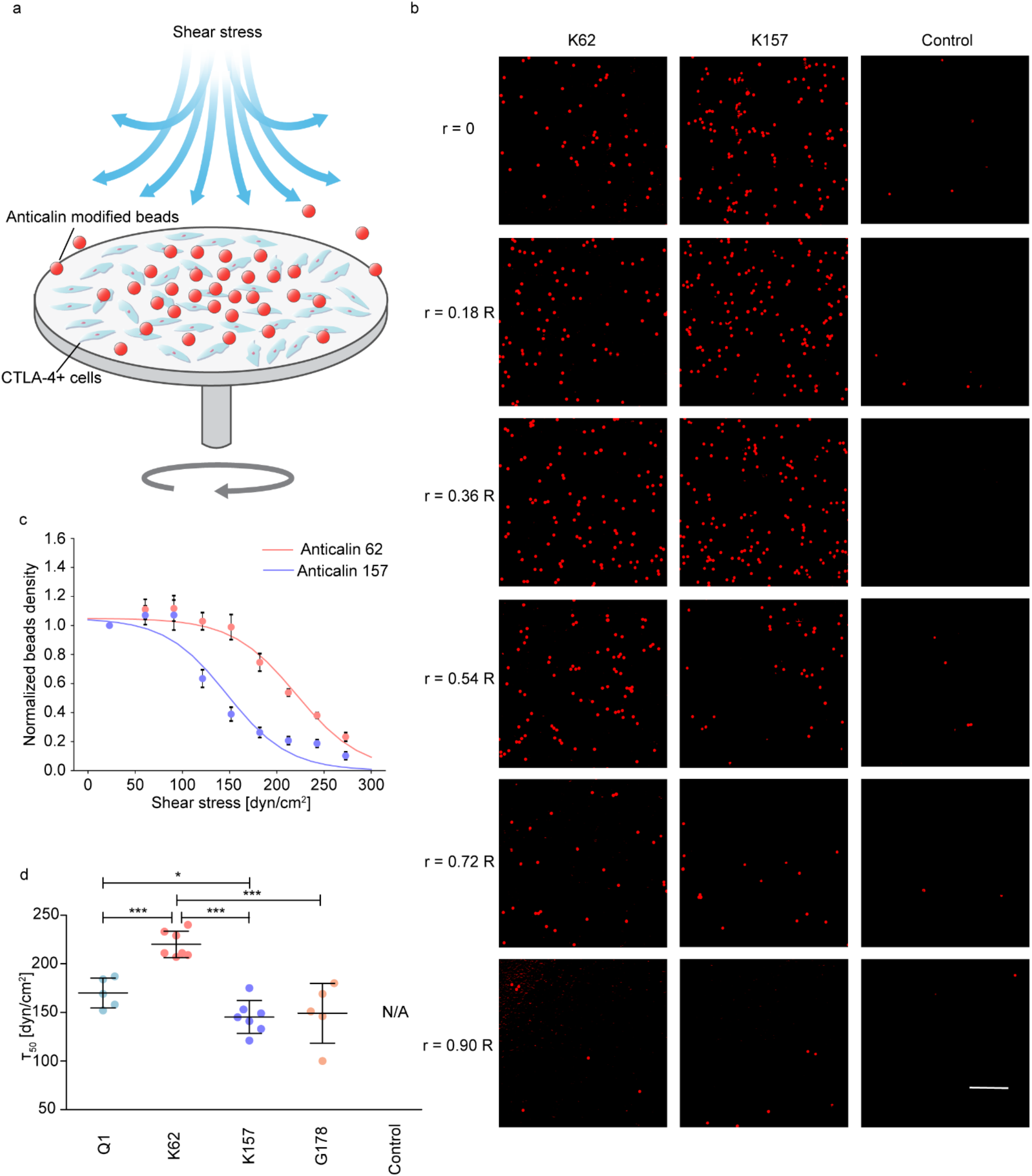
Quantitative measurement of bead adhesion to CTLA-4+ cells. **a**, Schematics of spinning disk experiments. Shear force is applied by the rotation of the substrate, and the binding strength between cells and Anticalin-modified beads is quantified by analyzing the spatial distribution of remaining beads. **b**, Fluorescence microscopy images of remaining beads on different regions with different distances to the center of the cover glass. Scale bar, 250 µm. **c**, Bead density versus shear stress for beads modified with Anticalin immobilized through anchor point K62 (red) and K157 (blue) residue. Data represent seven independent experiments. The experimental points for each Anticalin mutation were fitted with a sigmoid model. The midpoint extracted from the sigmoid fitting indicated the shear stress required to detach half the beads population (τ_50_). **d**, τ_50_ values of different bead samples where Anticalin was immobilized through different anchor points. Anticalin 1, cyan; Anticalin 62, red; Anticalin 157, purple; Anticalin 178, brown. Statistically significant differences are indicated as *P < 0.05, **P < 0.01, and ***P < 0.001.

We exposed cover glasses to shear stress (0-300 dyn/cm^2^) and imaged them with a fluorescence microscope. The beads lacking Anticalin modification served as a negative control. **Fig. 4b** shows a typical set of images from a spinning disk experiment. Beads lacking Anticalin modification were not retained on the cover glasses after spinning, demonstrating that non-specific binding was very weak under shear stress. In the group of beads modified with Anticalin attached through position K62 or K157, a substantial number of beads were observed in the areas of low shear stress (r = 0, r = 0.18 R, r = 0.36 R, where R is the cover glass radius). However, a notable decrease in bead density was observed at r = 0.54 R in the K157 anchor point group, while the K62 group experienced only a minor reduction. At positions further from the center on the cover glasses (r = 0.72 R, r = 0.90 R), the shear stress increased to levels that nearly removed all beads.

We quantified the bead density along the radius to create adhesion profile plots. These profiles were generated from multiple independent experiments and the data points were combined and globally fit to a sigmoidal function. This analysis determined the midpoint detachment shear stress (τ_50_)—the shear stress required to decrease the bead density to half of the maximum value at the center of the spinning disk. A higher τ_50_ indicates that higher shear stress is required to shear the beads off the cell surface, i.e., a greater adhesion strength. Notably, beads modified with Anticalin through the K62 anchor point demonstrated a significantly higher τ_50_ (220.0 ± 5.1 dyn/cm^2^) compared to those at the K157 anchor point (145.3 ± 6.4 dyn/cm^2^), as shown in **Fig. 4c**. We analyzed the τ_50_ values for beads modified at anchor points Q1, K62, K157, and G178 and plotted their respective adhesion profiles (**Figs. S9-12**). Statistically, the τ_50_ values for beads modified with Anticalin through position K62 were higher than those at other anchor points (**Fig. 4d**). Overall, results from the spinning disk experiments reveal that beads modified with Anticalin at the K62 anchor point exhibited the highest mechanical binding stability compared to those modified at the Q1, K157, and G178 anchor points. These findings are consistent with results from single-molecule force spectroscopy (SMFS) and rotary shaking experiments.

## Conclusion

To improve the binding strength of a binding scaffold to its antigen, protein engineers typically rely on genetic alterations (i.e. mutations) to the paratope, or other sequence variations that can change parameters such as hydrogen bonding with the antigen, charge-charge interactions, or hydrophobic character, and several others. Here we demonstrated an alternative concept for enhancing interaction strength under mechanical perturbation that we refer to as anchor point engineering. We showed how altering the immobilization point of a non-antibody scaffold Anticalin can be used to tune the mechanical stability of the complex with its target, CTLA-4. Anchor points were introduced into Anticalin using amber codon suppression and non-canonical azide incorporation. The azide anchor points were then derivatized with a mechanostable peptide handle for AFM-SMFS measurements. Single-molecule force measurements demonstrated how the various pulling geometries gave rise to differences in the energy landscape of the complex. We used multiparametric regression to discover that the distance of the pulling point from the CTLA-4 center of mass was significantly correlated with complex mechanostability. The generalizability of this finding remains to be tested in a wider range of protein-protein interaction systems. Bead adhesion assays performed on CTLA-4 positive cells under conditions that controlled for Anticalin avidity at the particle surface corroborated the results from the single-molecule analysis, and demonstrate adhesion enhancement when Anticalin was immobilized through the most mechanostable position (K62). These findings suggest a new approach for enhancing target binding strength in the context of particle-cell interactions under flow, and pave the way for engineering binding scaffolds that resist mechanical tension.

## Acknowledgments

This work was supported by the University of Basel, ETH Zurich and a Consolidator Grant (M822.00050) from the Swiss State Secretariat for Education, Research and Innovation (SERI) to MN.

## Data Availability

The data that support the findings of this study are openly available in zenodo at [*will be inserted prior to publication], reference number [*will be inserted prior to publication].

## Table of Contents Graphic

**Caption:**
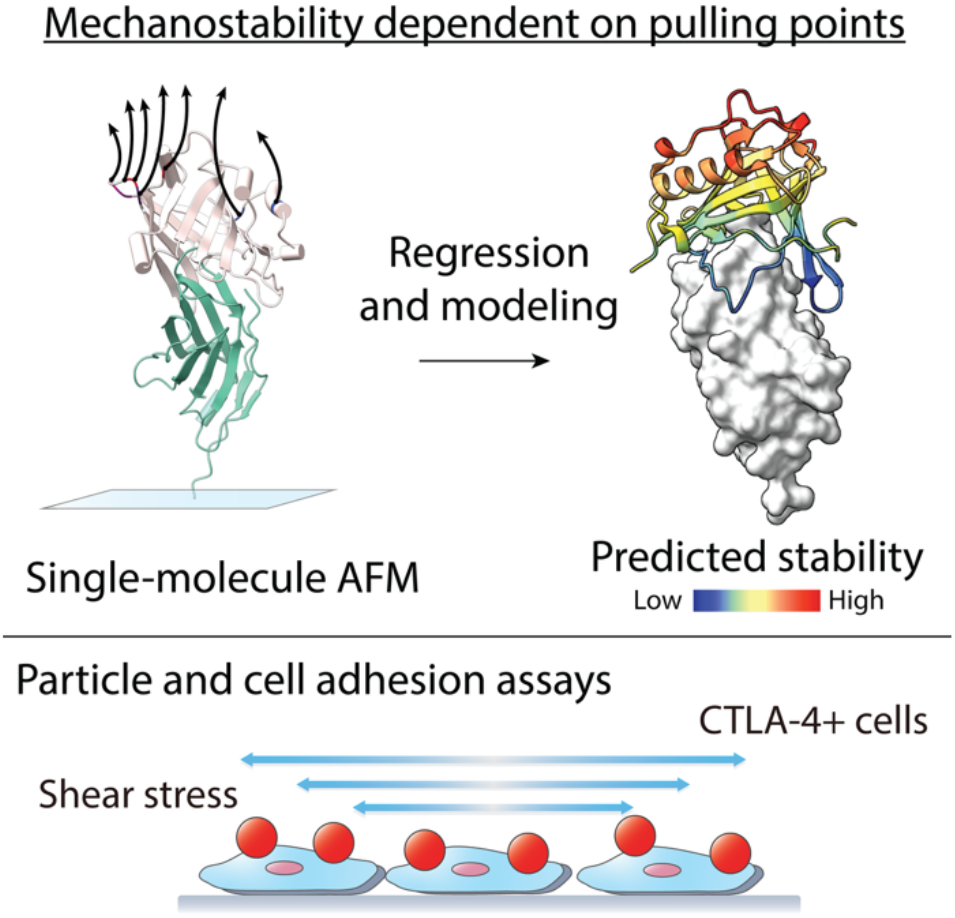
Anchor-point engineering modulates the mechanostability of targeting proteins. Clickable residues were placed at 17 sites on a CTLA-4-binding Anticalin, and probed by AFM and particle-based adhesion assays. Pulling from K62 located farthest from CTLA-4’s center of mass generated the strongest molecular complexes and highest particle adhesion. This work introduces a simple design rule for functionalizing particles to adhere to specific cell types and resist mechanical shear.

