## Supplementary Information for "Engineering Mechanostable Anticalin Scaffolds to Enhance Particle Adhesion and Targeting of CTLA-4 under Shear Stress"

Yang Sun[1,2†], Jiajun Li[1,2†], Rosario Vanella[1,2], Haipei Liu[1,2], Zhaowei Liu[1,2,3], and Michael A. Nash[1,2*]
[1] Institute of Physical Chemistry, Department of Chemistry, University of Basel, 4058 Basel, Switzerland
[2] Department of Biosystems Science & Engineering, ETH Zurich, 4056 Basel, Switzerland
[3] Present address: Department of Physics, Oxford University, OX1 3QU United Kingdom

### Methods and Materials

#### Expression and purification of anticalin with Amber suppression

Internal anchor points were introduced to different residues of anticalin constructs by utilizing an amber suppression system. The original sequence of the wild-type anticalin sequence was inserted into a pET28a vector and the codon corresponding to the amino acid at the designated anchor point was mutated to an amber codon (TAG). NiCo21(DE3) competent cells (New England Biolabs, Ipswich, MA, USA) were co-transformed with both the expression vector and the pEVOL-pAzF plasmid (addgene #31186). Cells were cultivated in LB medium supplemented with 50 µg/mL of ampicillin, 25 µg/mL of chloramphenicol, and 0.2 mg/ml of p-azido-l-phenylalanine (pAzF) at a temperature of 37 °C until the optical density (OD) was approximately 0.6. Subsequently, the culture was further incubated at 37 °C for 1 h with the addition of 0.2 mg/mL of arabinose, followed by an overnight incubation at 20 °C with 1 mM of isopropyl β-D-1-thiogalactopyranoside (IPTG). After incubation, the cells were collected by centrifugation, lysed through sonication, and the lysate was loaded to the His-trap column (GE Healthcare, IL, USA). The column was washed with PBS buffer (137 mM NaCl, 2.7 mM KCl, 10 mM $Na_2HPO_4$ and 2 mM $KH_2PO_4$, pH = 7.4) containing 20 mM imidazole and further eluted with PBS buffer supplemented with 500 mM imidazole. The eluate underwent further purification with a Superdex Increase 200 10/300 GL size-exclusion column (SEC) provided by GE Healthcare.

#### Conjugation of Fgβ peptide and anticalin

A 1.5x molar excess of Fgβ-StrepTag-DBCO peptide (JPT Peptide Technologies GmbH, Berlin, Germany) was mixed with AzF-incorporated anticalin. The resulting mixture underwent incubation at room temperature with shaking for 1 hour, followed by an overnight incubation at 4 °C. Subsequently, the reaction mixture was loaded to a SEC to eliminate any unbound peptide. The protein obtained through SEC purification was then applied to a Strep-Trap column (GE Healthcare) and eluted using PBS buffer containing 2.5 mM desthiobiotin to eliminate any unreacted anticalin.

#### Expression and purification of CTLA-4-ddFLN4-ELP-ybbr

SHuffle® T7 competent cells (New England Biolabs, Ipswich, MA, USA) were transformed with the pET28a vector carrying CTLA-4-ddFLN4-ELP-ybbr. Cells were cultured in TB medium supplemented with 50 µg/mL of kanamycin at 30 °C until OD was approximately 0.6. The expression of CTLA-4-ddFLN4-ELP-ybbr was induced by adding 0.4 mM of IPTG followed by an overnight incubation at 16 °C. The CTLA-4-ddFLN4-ELP-ybbr protein was purified using the same procedure as the Fgβ-anticalin construct.

**Surface chemistry for AFM measurements**

Cover glasses and Biolever mini (Bruker, Billerica, MA, USA) AFM cantilevers were amino-silanized using (3-aminopropyl)-dimethyl-ethoxysilane (APDMES, ABCR GmbH, Karlsruhe, Germany). The silanized surfaces were then subjected to incubation in 10 mg/mL of sulfosuccinimidyl 4-(Nmaleimidomethyl) cyclohexane-1-carboxylate (sulfo-SMCC, Thermo Fisher Scientific, Waltham, MA, USA) at room temperature for 30 min, followed by another incubation in 200 μM of coenzyme A (CoA, Sigma-Aldrich, St. Louis, MO, USA) at room temperature for 2 h. The cover glasses and cantilevers were thoroughly washed with dd$H_2O$ after each incubation step. The CoA-coated cantilevers and cover glasses were incubated in a CoA-ybbr reaction mixture, comprising 40 μM of ybbr-tagged protein, 5 μM of Sfp (phosphopantetheinyl transferase) enzyme and 20 mM of $MgCl_2$, and incubated at room temperature for 2 hours. Subsequently they were washed with PBS buffer and stored in PBS until further use.

**AFM measurements**

AFM-SMFS (Atomic Force Microscopy-based single molecule force spectroscopy) measurements were performed using a Force Robot AFM (JPK instruments, Berlin, Germany). The cantilever spring constants (ranging from 0.02 N/m to 0.14 N/m) and detector sensitivity were calibrated using the contact-free method. The cantilever and cover glasses were immersed in PBS buffer. 1 μM of freely diffusing Fgβ-conjugated anticalin was added to the measurement buffer. In the experiments, the cantilever was brought into proximity to the glass surface, dwelled for 200 ms and then retracted at a constant speed within the range of 100 nm/s to 3200 nm/s (100 nm/s, 400 nm/s, 800 nm/s and 3200 nm/s). After each approach-retraction cycle, the x-y position of the stage was displaced by 100 nm to probe a new molecule on the surface.

**AFM data analysis**

During the measurement, the collected force curves were initially subjected to real-time analysis and filtering using software integrated into the AFM instrument. The criteria applied were aiming to exclude plain force traces and non-specific binding to the glass substrate. Specifically, the maximum adhesion force of the retraction segment with a minimum extension of 10 nm needed to exceed 30 pN. Following the measurement, the selected force curves exhibiting positive force signals underwent further processing and analysis through contour length transformation. Since Sfp surface chemistry includes a fingerprint domain of ddFLN4 for CTLA4 and anticalin, respectively. Therefore, single molecular force traces were identified by searching for contour length increments matching the fingerprint domain of 2 × ddFLN4 (≈ 2 × 32 nm). For each force-distance trace, the loading rate at the point of rupture was extracted by applying a line fit to the force versus time trace in the immediate vicinity before the rupture peak. The loading rate was determined from the slope of the fit. For each pulling speed, the distribution of loading rates was fitted with Kernel Density Estimation (KDE), and the peak loading rate was obtained from the fitted curve. The rupture forces measured at different pulling speeds were plotted in histograms and the rupture force distribution p(F) was fitted with Bell-Evans model to extract the most probable rupture force. Both Bell-Evans (BE) model and Dudko-Hummer-Szabo (DHS) model were used to calculate the energy barrier parameters including the distance to the energy barrier $\Delta x$ (BE), the off rate at zero force $k_0$ (BE), $\Delta x$ (DHS), the height of the energy barrier in the absence of force $\Delta G$ (DHS), and the zero-force life time $\tau_0$ (DHS) as described before. [25]

**Statistical analysis**

Quantitative data are expressed as the mean ± the standard error of the mean (S.E.M.) derived from a minimum of three independent experiments. Statistical analysis was conducted using GraphPad Prism software. The p-value was computed using the unpaired t test with Welch's correction with a two-tailed model and a confidence level of 95%. A p-value of 0.05 was considered statistically significant.

**Modification of polybeads with anticalin**

Anticalin with amber mutation at sites Q1, K62, K157, and G178 were selected for further cellular-level investigation. 400 μL of polybeads (Φ20 μm, 3D-DBCO, PolyAn Plex Red4, 0.5% solids (polybeads, PolyAn, Berlin, Germany) stock solution was washed by PBS 3 times and resuspended in 100 μL of PBS. Subsequently, for the equal avidity group, 0.175/1.375/1.25/4 μM of anticalin Q1/K62/K157/G178 was separately added to the polybeads solution. While for the maximal avidity group, 4 μM of anticalin Q1/K62/K157/G178 was separately added to the polybeads solution. And then the total volume was made up by PBS to 1 mL. The mixture was incubated at 4 °C overnight. Following incubation, the polybeads were washed with PBS twice. They were then blocked at room temperature with 1% BSA for 1 hour. After blocking, the polybeads were washed with PBS and incubated with 1 μg/mL of the primary antibody (6x-His Tag, MA1-21315, Invitrogen, USA) for 1 hour at room temperature. Subsequent to washing by centrifugation, the polybeads were further incubated with 2 μg/mL of the secondary antibody (Goat anti-Rabbit IgG, Alexa Fluor Plus 488, A32731, Invitrogen, USA) for 1 hour at room temperature and washed again by centrifugation. Polybeads contain red fluorescent signals, and successful conjugation of secondary antibodies to anticalin modified on the surface of polybeads introduced additional green fluorescent signals, which were assessed by the median distribution of the green fluorescent intensity measured in flow cytometry. The amount of anticalin with different anchor points used in modification was adjusted to ensure similar anticalin modification level on the polybeads.

**Micro-shaking cell binding experiments**

Human CTLA-4+ CHO-K1 cells (MyBioSource, Inc., San Diego, CA, USA) at passages 4 to 10 (P4-P10) were cultured in DMEM medium (w/ L-Glutamine, 4.5 g/L Glucose and Sodium Pyruvate), supplemented with 10% FBS and 1% Pen/Strep, plus 10 μg/ml of Blasticidin and passaged every week in a 1:20 ratio. CTLA-4+ CHO-K1 cells were seeded onto 48-well plates and allowed to adhere and grow for 1 week. 25 μL of polybeads stock solution modified with the equivalent/maximized avidity of anticalin at different anchor points were added to each well. External forces can be exerted to the system by placing the cell culture plate on a Microplate Vortex Mixer (OHAUS, Parsippany, New Jersey, USA). After incubating for 1 hour at 300/600/900 RPM, the culture media was removed and cells were washed with D-PBS twice, followed by a subsequent shaking culture at 300/600/900 RPM for another 1 hour. Next, cells were washed with HBSS (Hank's Balanced Salt Solution) to remove unconjugated polybeads. Subsequently, cells were stained in a 2 μM of Calcein, AM solution in HBSS at 37 °C for 30 minutes, and then the culture medium was used to replace the staining solution. The binding efficiency of polybeads against CTLA-4+ cells was evaluated by normalizing the number of conjugated beads by the number of live cells obtained from fluorescence microscopy.

**Spinning disk cell binding assay**

The spinning disk apparatus was built in lab as reported previously. CTLA-4+ CHO-K1 cells were seeded on the surface of aminosilanized coverglasses in 6-well plates and allowed to adhere and grow for 1 week in culture medium. 200 μL of polybeads stock solution modified with the equivalent avidity of anticalin at different anchor points were added to each well. After incubating for 1 hour at 600 RPM, the cover glasses were washed subsequently with HBSS (Hank's Balanced Salt Solution, Invitrogen, Cat. No. 14025-092) to remove unconjugated polybeads. Cells were stained in 2 μM of Calcein, AM (Invitrogen, Cat. No. C1430) in HBSS at 37 °C for 30 minutes. The cover glasses were mounted on the spinning disk device, secured by vacuum suction, and immersed in culture medium at room temperature. The spinning routine consisted of a 2.5 s acceleration ramp, 5 min steady spinning at 2500 rpm value, and 2.5 s deceleration. During the binding assay, the cover glasses were maintained at a height of 25 mm from the bottom of the chamber to minimize turbulence and not disrupt the laminar boundary layer. The shear stress (τ; Pa) at any point on the surface of the cover glass varies linearly with radial distance and is described as in the previous paper.[36] After spinning, the cover glass was raster scanned and imaged frame by frame at 10× magnification on an Olympus IX81 microscope (~500 individual images automatically stitched together with CellSens software (version 1.16; Olympus, Tokyo, Japan)) and saved in TIFF format. The image analysis was performed in ImageJ 1.54f. Polybeads were segmented by gray-value thresholding. The coordinates of polybeads were acquired as csv files by "Analyze Particles" function in ImageJ. Further data analysis was performed in Python script. The fraction of adherent polybeads at different positions on the disk was calculated by normalizing the density of polybeads at each section of the disk with that at the center of the disk, where the shear forces are close to zero. We plotted the detachment profiles and fitted them with a sigmoid probabilistic model: $f=a/(1+\exp[b(\tau-\tau 50)])$, where τ50 is the shear stress at which 50% of polybeads remain adherent. This value was used as a measure of mean adhesion strength for comparison of polybeads populations.

**Microscale thermophoresis (MST) measurements**

Anticalin with amber mutation at sites Q1, K62, K157, and G178 were labeled with the RED-NHS 2nd generation dye solution (NanoTemper Technologies GmbH, München, Germany) by mixing 90 μL of 10 μM anticalin with 10 μL of the 300 μM dye solution and incubate at room temperature in the dark for 30 minutes. The labeled proteins were purified from unreacted free dye using B-column, which was equilibrated with PBS. The titration samples were prepared by mixing 80 nM of labelled anticalin with a series of unlabelled CTLA-4 with different concentrations ranging from 488 pM to 32 μM in a 1:1 ratio. The microscale thermophoresis (MST) traces of each titration sample were measured using a Nanotemper Monolith NT.115 (NanoTemper Technologies GmbH, Munich, Germany). The temperature-related fluorescence intensity change was recorded for each sample and plotted against the concentration of CTLA-4 to derive the dissociation constant between anticalin and CTLA-4. $F_{norm}$ is defined as the ratio between the average fluorescence in the hot region and the cold region in the MST time trace. The cold and hot regions have been defined as the 1 s region just before the T(temperature)-Jump and the 5 s region after the T-Jump. The dissociation constant $K_D$ was fit from the binding curve using the 1:1 binding model as described in the literature.[37]

**Multi-parameter linear regression fitting**

The multi-parameter linear regression was performed with a Python script. The features including distance of the anchor point to the CTLA4 mass center, the hydropathy of the mutated amino acid, electronic charge index of the mutated amino acid, and solvent accessible surface area of the mutated amino acid were scaled by its maximum absolute value. The linear regression was done by Linear Regression function in the Scikit-Learn. linear_model module. The importance of the features was evaluated by F test using the SelectKBest function in the Scikit-Learn. feature_selection module.

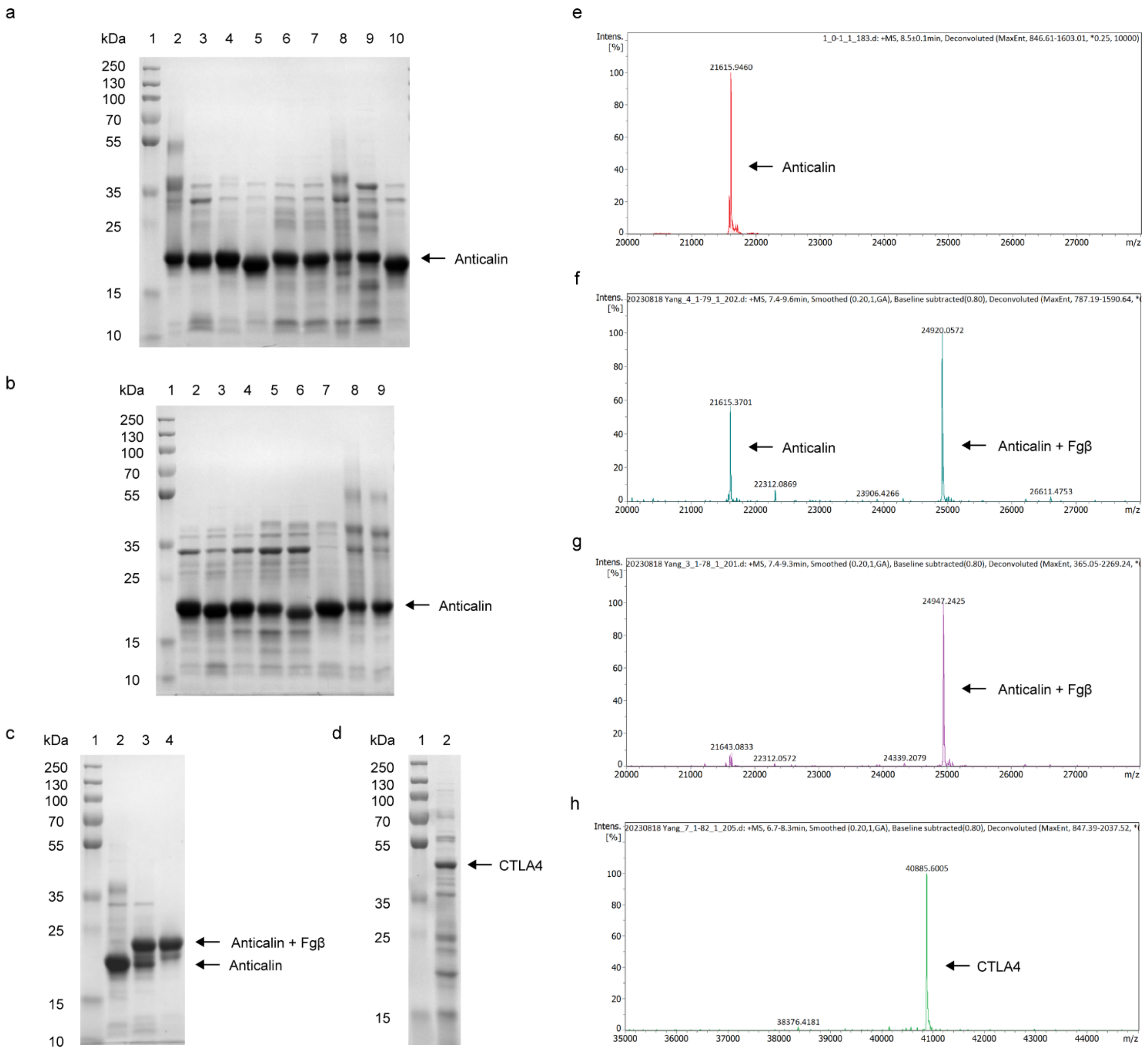


**Figure S1 |** Purification of anticalin and CTLA-4, and successful conjugation of Fgβ to anticalin demonstrated by SDS-PAGE and MS. **a**, SDS-PAGE gel of marker (lane 1) and anticalin with Amber suppression at Q1 (lane 2), Y32 (lane 3), I55 (lane 4), E57 (lane 5), K59 (lane 6), E60 (lane 7), K62 (lane 8), S63 (lane 9), S71 (lane 10). **b**, SDS-PAGE gel of marker (lane 1) and anticalin with Amber suppression at Y78 (lane 2), S87 (lane 3), T93 (lane 4), L107 (lane 5), K124 (lane 6), E143 (lane 7), K157 (lane 8), G178 (lane 9). **c**, SDS-PAGE gel of marker (lane 1) and anticalin with Amber suppression at S87 purified after SEC column (lane 2), conjugated with Fgβ peptide and purified after SEC column (lane 3), and then purified with Strep-Trap column (lane 4). Successful conjugation of the peptide increased the protein molecular weight by ~3 kD. **d**, SDS-PAGE gel of marker (lane 1) and CTLA-4 purified after SEC column. **e**, Measurements on anticalin S87AzF mutant before conjugation with Fgβ-StrepTag-DBCO, **f**, conjugated anticalin after SEC column purification, **g**, purified further after Strep-Trap column. The theoretical molecular weight of S87AzF mutant is 21,615 Da. After conjugation with the Fgβ peptide, the molecular weight increased by 3,305 Da, exactly matching the molecular weight of the synthetic Fgβ-StrepTag-DBCO peptide. **h**, CTLA-4 after SEC column purification.

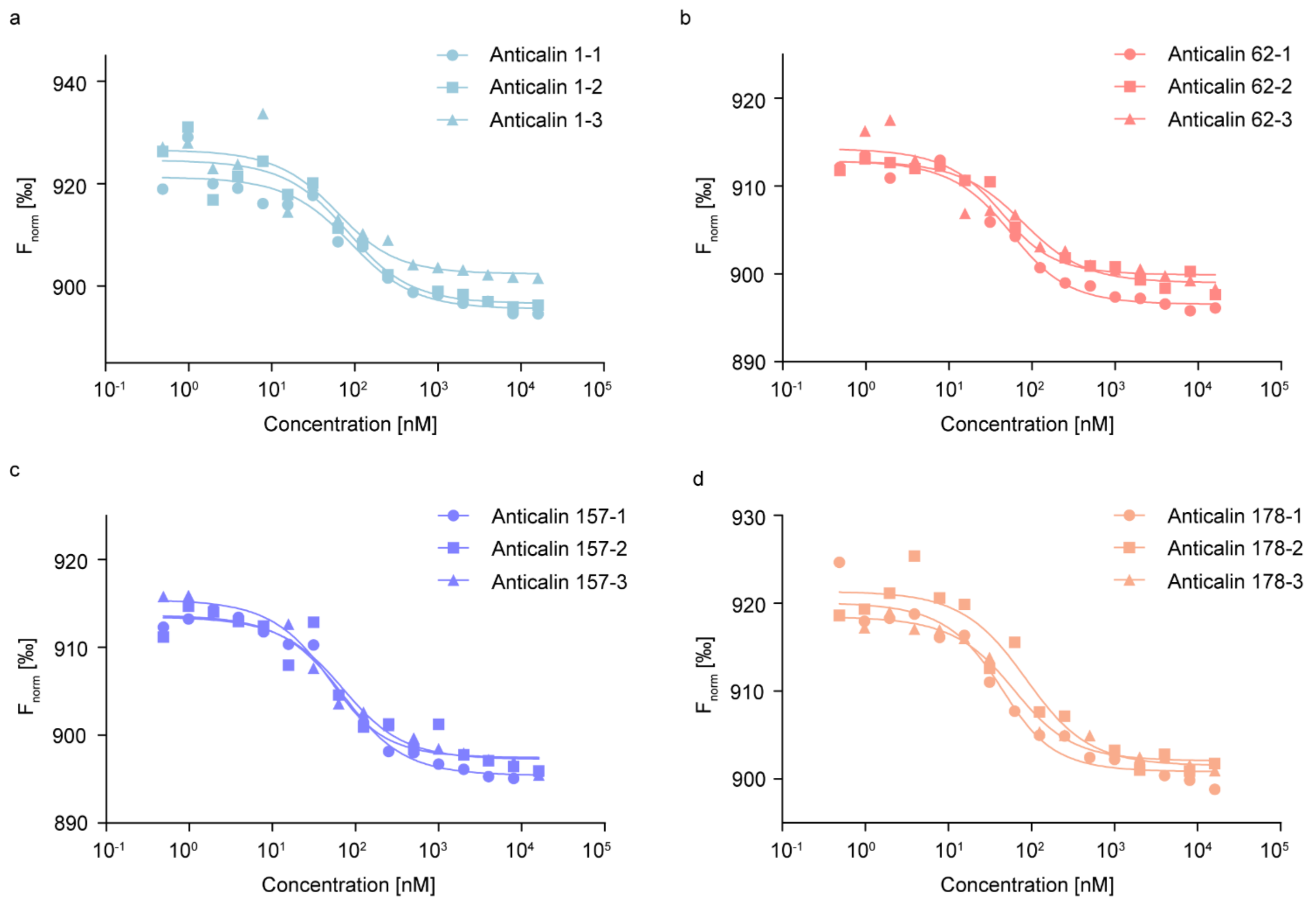


**Figure S2 |** Equilibrium binding affinity by thermophoresis. MST binding curves of anticalin with Anchor points at **a**, Q1, **b**, K62, **c**, K157 and **d**, G178. The CTLA-4 was titrated against fluorescent labelled 40 nM anticalin. The normalized thermophoresis value $F_{norm}$ was plotted against the CTLA-4 concentration. Data of three independent experiments were shown. Dissociation constants ($K_D$) were fitted with a 1-1 binding model. Mean values of $K_D$ ± standard deviation for CTLA-4 binding with anticalin from three independent experiments were shown in **Table S3**.

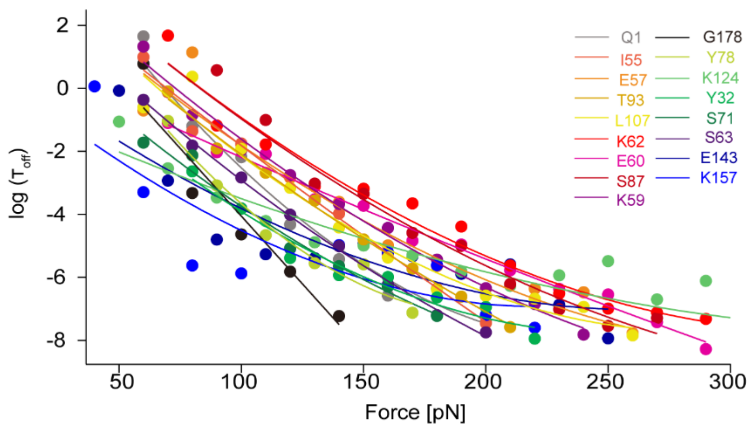


**Figure S3 |** Force-dependent lifetime of the (Anticalin:CTLA-4) complex. The force-dependent lifetime was plotted against force and fit using the DHS model to extract distance to the energy barrier $\Delta x$ (DHS), the height of the energy barrier in the absence of force $\Delta G$ (DHS), and the zero-force life time $\tau_0$ (DHS).

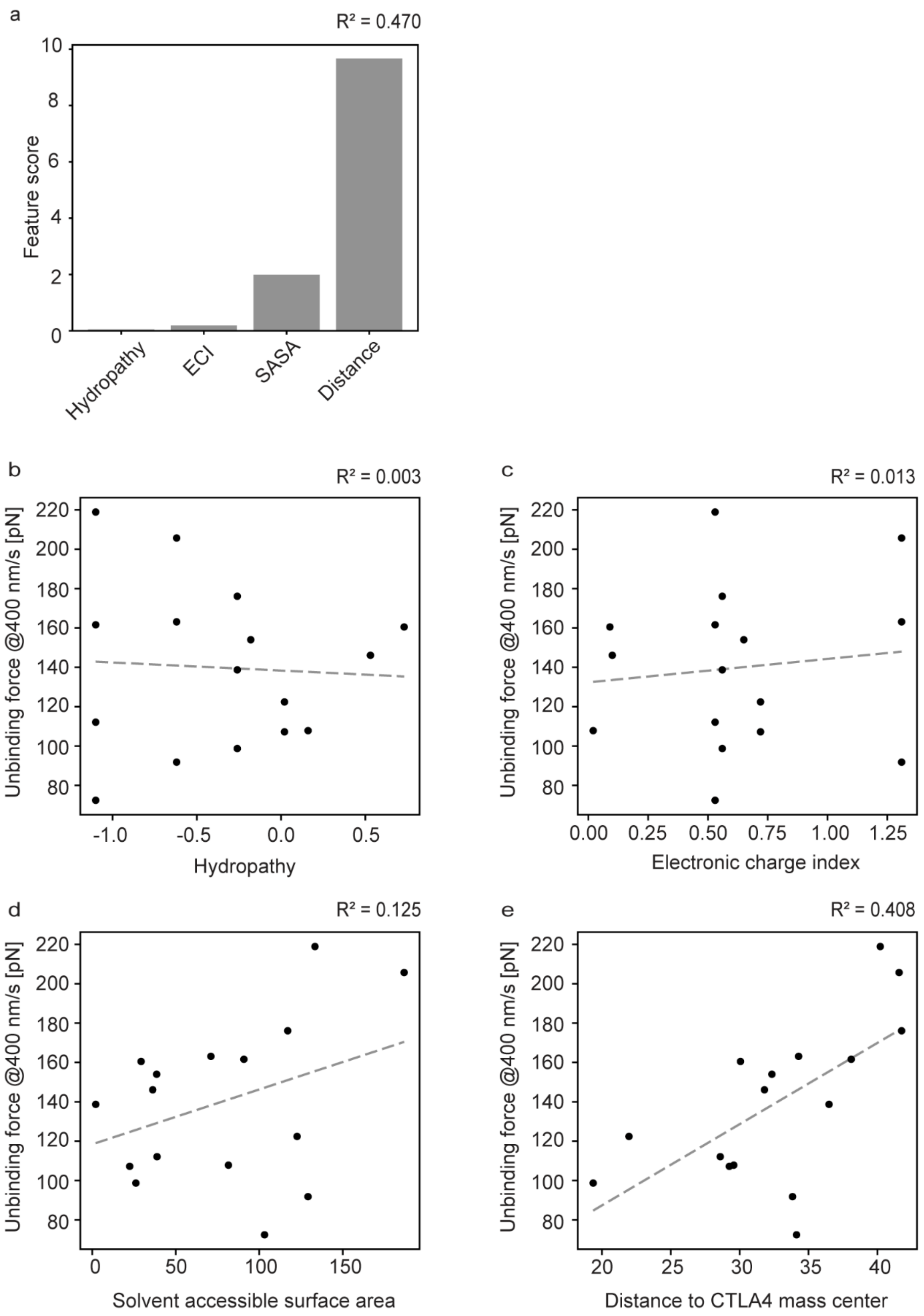


**Figure S4 |** Physicochemical properties of all anticalins. **a**, The bar graph of the feature score of different Physicochemical properties. **b**, Linear regression of the hydropathy against the unbinding force at 400 nm/s. ($R^2$=0.003). **c**, Linear regression of the electronic charge index against the unbinding force at 400 nm/s. ($R^2$=0.013). **d**, Linear regression of the solvent accessible surface area against the unbinding force at 400 nm/s. ($R^2$=0.125). **e**, Linear regression of the distance to CTLA-4 mass center against the unbinding force at 400 nm/s. ($R^2$=0.408).

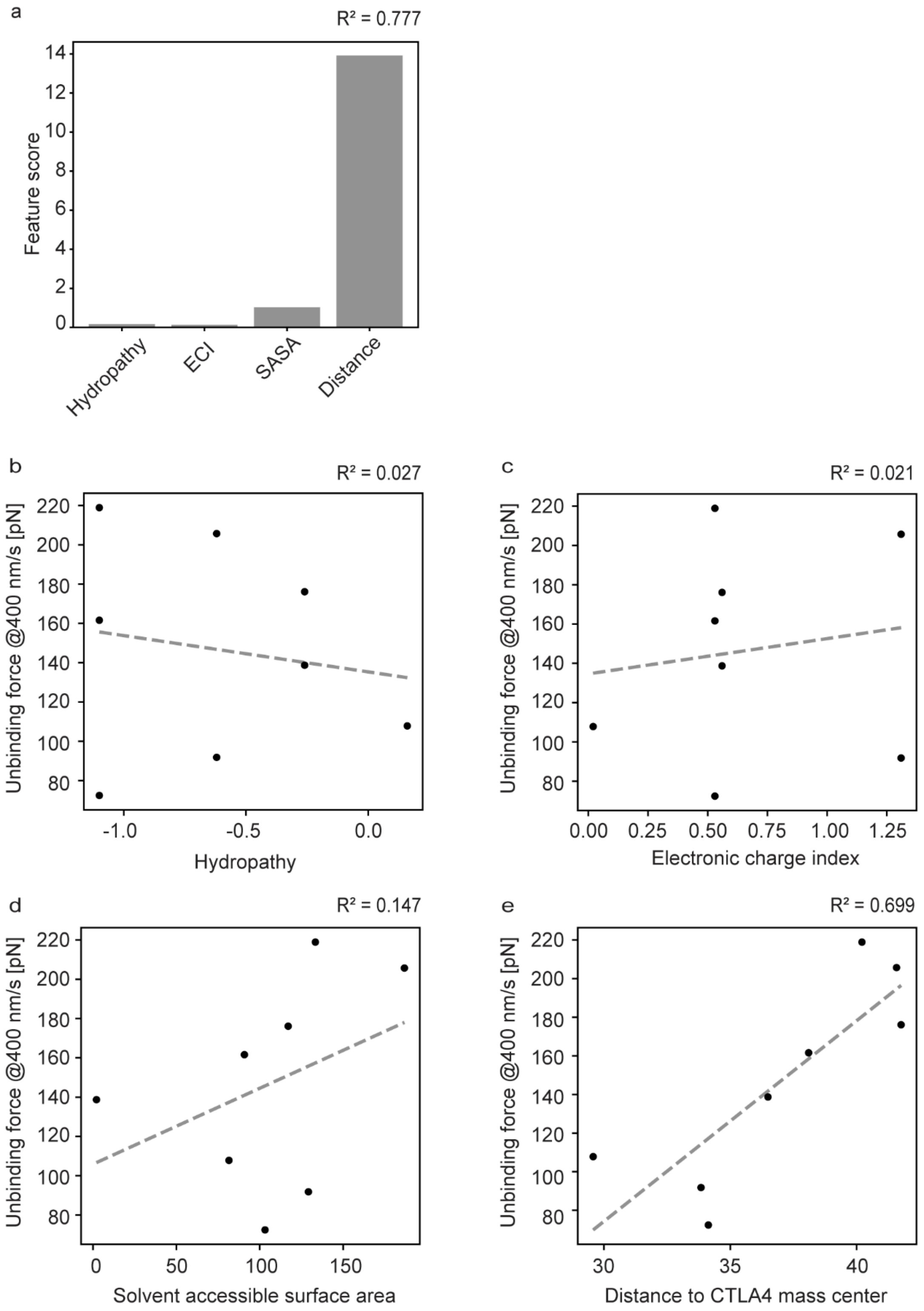


**Figure S5 |** Physicochemical properties of anticalins with anchor points at loops and alpha helices. **a**, The bar graph of the feature score of different Physicochemical properties. **b**, Linear regression of the hydropathy against the unbinding force at 400 nm/s. ($R^2$=0.027). **c**, Linear regression of the electronic charge index against the unbinding force at 400 nm/s. ($R^2$=0.021). **d**, Linear regression of the solvent accessible surface area against the unbinding force at 400 nm/s. ($R^2$=0.147). **e**, Linear regression of the distance to CTLA-4 mass center against the unbinding force at 400 nm/s. ($R^2$=0.699).

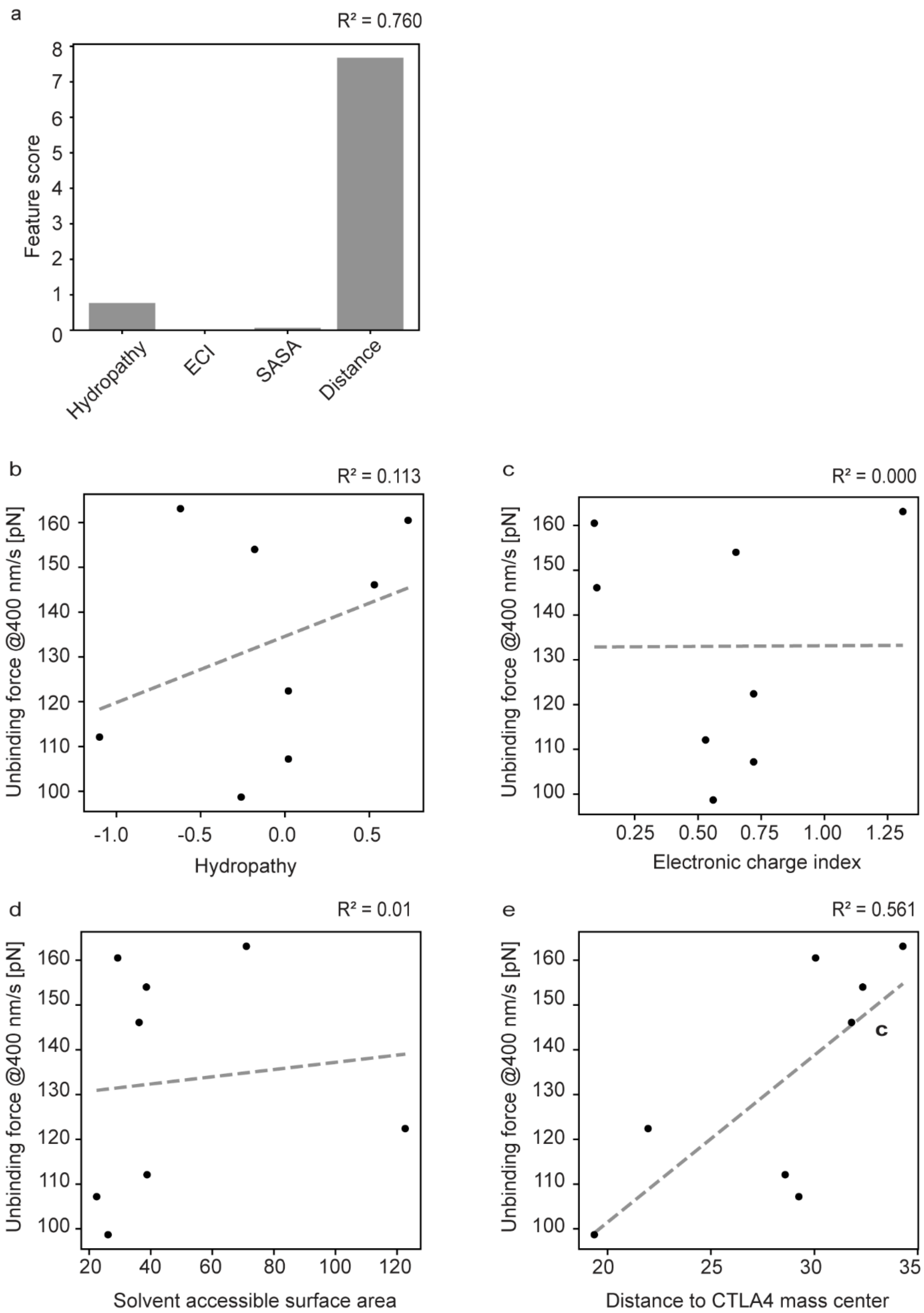


**Figure S6 |** Physicochemical properties of anticalins with anchor points at beta strands. **a**, The bar graph of the feature score of different Physicochemical properties. **b**, Linear regression of the hydropathy against the unbinding force at 400 nm/s. ($R^2$=0.113). **c**, Linear regression of the electronic charge index against the unbinding force at 400 nm/s. ($R^2$=0.000). **d**, Linear regression of the solvent accessible surface area against the unbinding force at 400 nm/s. ($R^2$=0.01). **e**, Linear regression of the distance to CTLA-4 mass center against the unbinding force at 400 nm/s. ($R^2$=0.561).

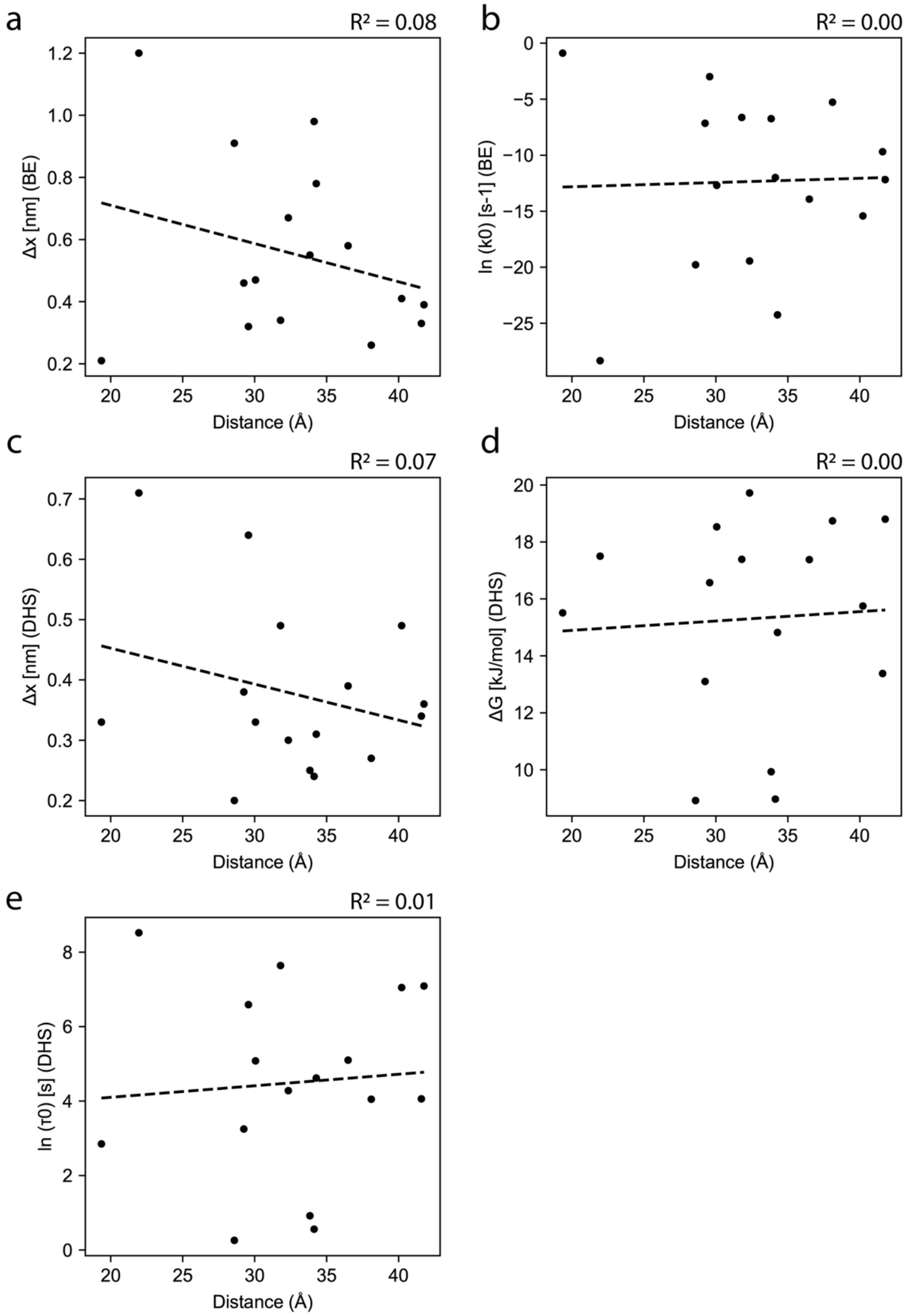


**Figure S7 |** Fitted energy landscape parameters are uncorrelated with distance between the anchor point and CTLA-4 center of mass. In each graph, a. linear regression is performed between an energy landscape parameters fitted from the loading rate-dependent SMFS data and the distance between the anchor point and the CTLA-4 center of mass. No significant correlations were found **a**, Fitted Δx (Bell-Evans) vs. distance. **b**, Fitted Ln($k_{off}$) (Bell-Evans) vs. distance. **c**, Fitted Δx (DHS) vs. distance. **d**, Fitted ΔG (DHS) vs. distance. **e**, Fitted Ln(τ**)** (DHS) vs. distance where τ is the bond lifetime.

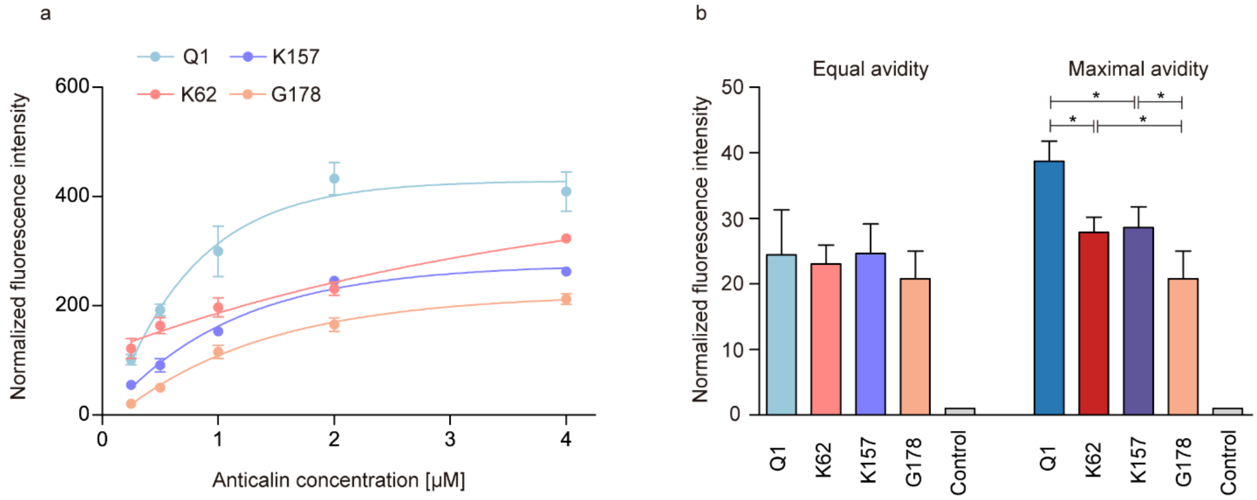


**Figure S8 |** Characterization of anticalin modified on the surface of polybeads. **a**, Fluorescence intensity of anticalin with different concentrations of Q1/K62/K157/G178 anchor points modified on the surface of polybeads. **b**, Modification of anticalin with Q1/K62/K157/G178 anchor points at equal avidity/maximal avidity.

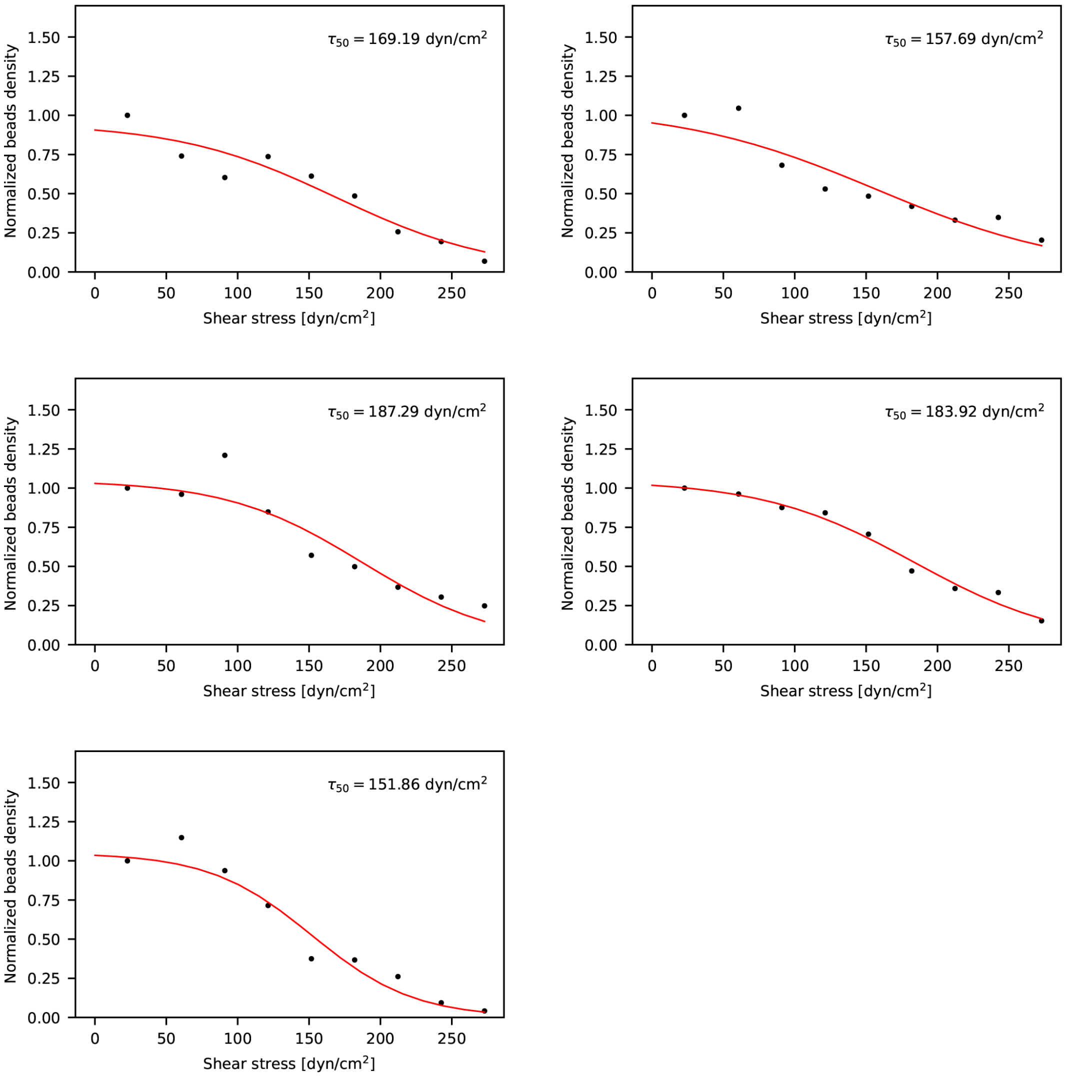


**Figure S9 |** Beads density versus shear stress plots for beads modified with anticalin with anchor points at Q1 residue. Each plot shows data from one of the five replicates. $T_{50}$ was calculated by fitting the experimental points for each beads' population versus shear stress.

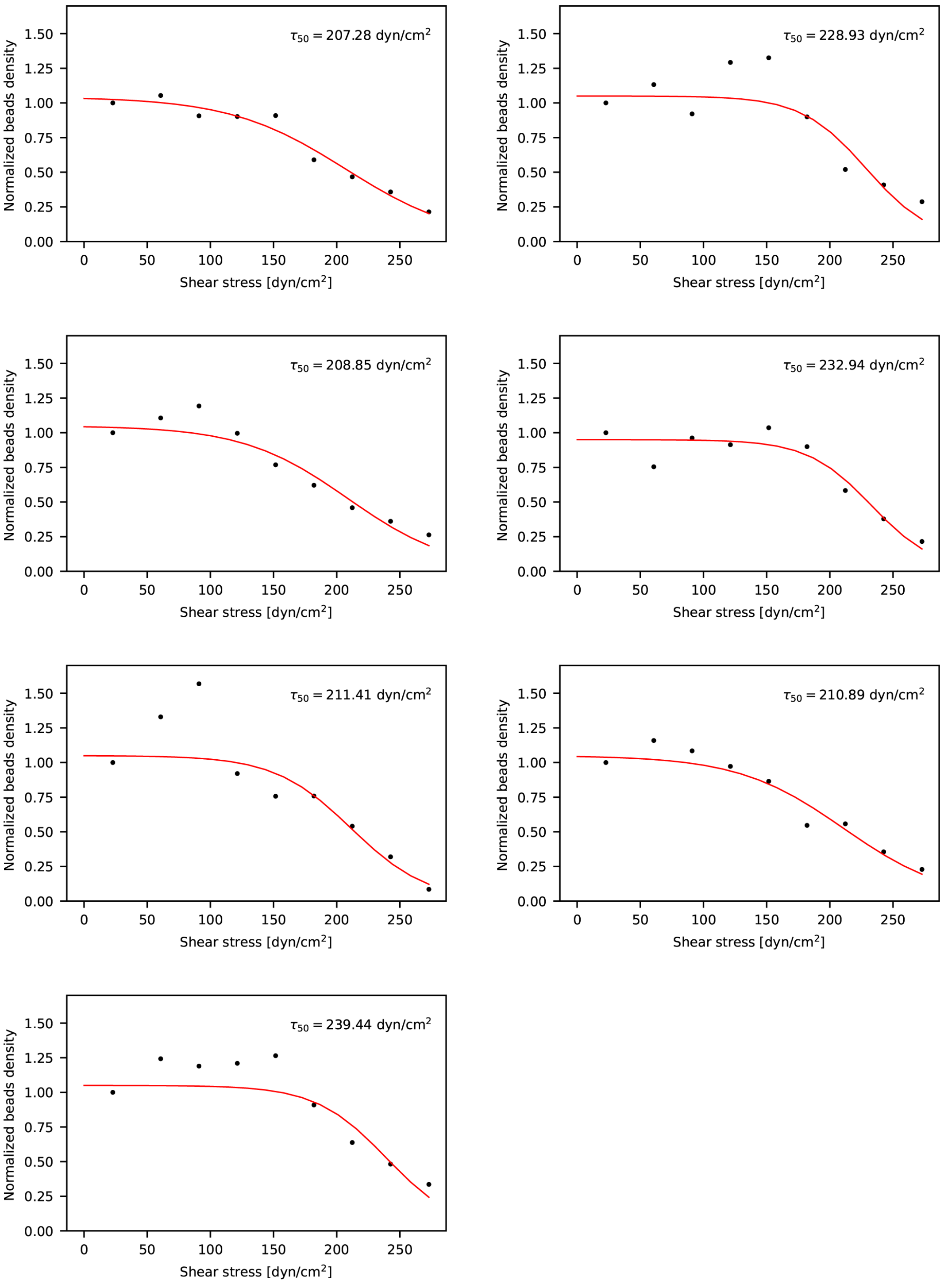


**Figure S10 |** Beads density versus shear stress plots for beads modified with anticalin with anchor points at K62 residue. Each plot shows data from one of the seven replicates. $T_{50}$ was calculated by fitting the experimental points for each beads' population versus shear stress.

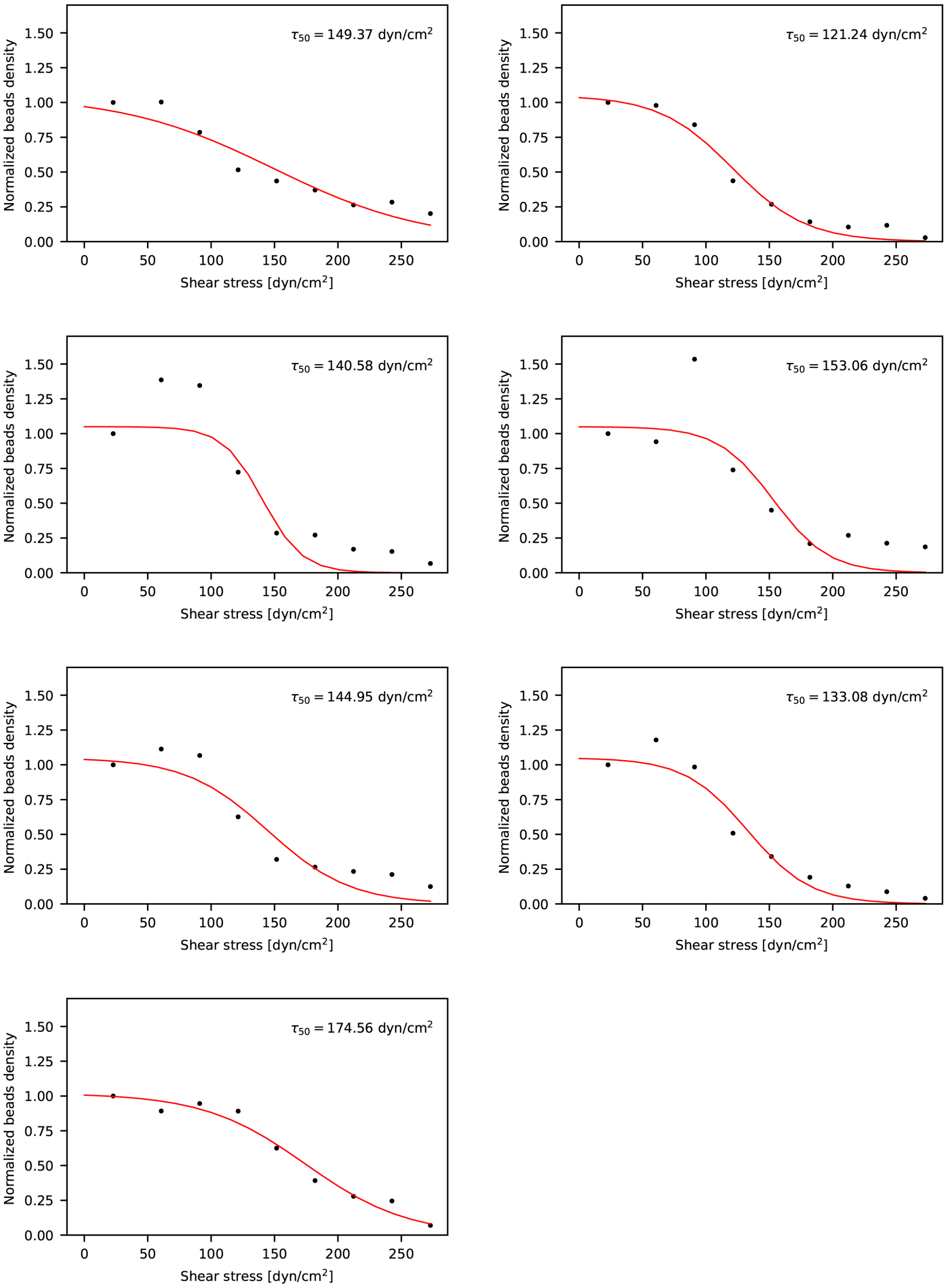


**Figure S11 |** Beads density versus shear stress plots for beads modified with anticalin with anchor points at K157 residue. Each plot shows data from one of the seven replicates. $T_{50}$ was calculated by fitting the experimental points for each beads' population versus shear stress.

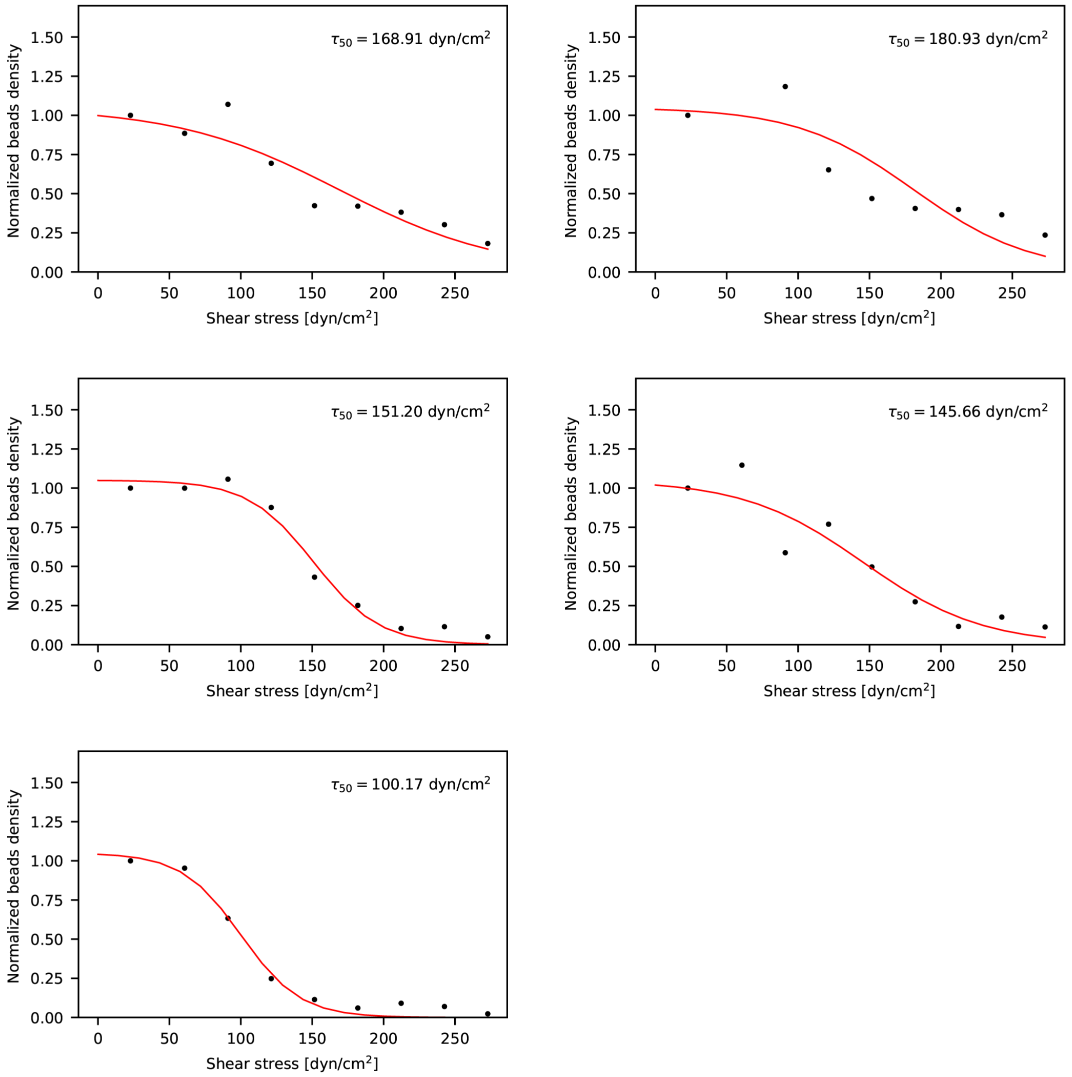


**Figure S12 |** Beads density versus shear stress plots for beads modified with anticalin with anchor points at G178 residue. Each plot shows data from one of the five replicates. $T_{50}$ was calculated by fitting the experimental points for each beads' population versus shear stress.

| Loop | Sequence | | Anchor points |
|---|---|---|---|
| L2 | 59-63 | KEDKS | K59 E60 K62 S63 |
| L4 | 86-90 | GSQPG | S87 |
| L8 | 140-144 | RTKEL | E143 |
| N/C ends | Sequence | | Anchor points |
| N | 1 | Q | Q1 |
| C | 178 | G | G178 |
| β strand | Sequence | | Anchor points |
| S1 | 29-38 | GKWYVVGLAG | Y32 |
| S2 | 53-58 | ATIYEL | I55 E57 |
| S3 | 64-72 | YNVTSVISS | S71 |
| S4 | 75-85 | KCFYTIATFVP | Y78 |
| S5 | 91-94 | EFTL | T93 |
| S6 | 105-113 | SYLVRVVST | L107 |
| S7 | 118-127 | YAVVFFKLAE | K124 |
| S8 | 130-139 | AEFFAITIYG | E131 (poor expression) |
| α helix | Sequence | | Anchor points |
| A3 | 145-159 | ASELKENFIRFSKSL | K157 |

**Table S1.** Anchor points design in anticalin.

| | Q1 | Y32 | I55 | E57 | K59 | E60 | K62 | S63 | S71 | Y78 | S87 | T93 | L107 | K124 | E143 | K157 | G178 |
|---|---|---|---|---|---|---|---|---|---|---|---|---|---|---|---|---|---|
| Q1 | | no | *** | *** | *** | *** | *** | ** | no | * | *** | *** | *** | no | *** | *** | no |
| Y32 | no | | ** | *** | *** | *** | *** | *** | no | *** | *** | *** | *** | no | no | *** | no |
| I55 | *** | ** | | *** | *** | *** | *** | *** | no | *** | *** | *** | *** | *** | no | * | *** |
| E57 | *** | *** | *** | | no | no | *** | *** | * | ** | no | no | no | *** | *** | *** | *** |
| K59 | *** | *** | *** | no | | * | *** | *** | * | *** | * | no | no | *** | *** | *** | *** |
| E60 | *** | *** | *** | no | * | | * | *** | ** | *** | no | * | ** | *** | *** | *** | *** |
| K62 | *** | *** | *** | *** | *** | * | | *** | *** | *** | ** | *** | *** | *** | *** | *** | *** |
| S63 | ** | *** | *** | *** | *** | *** | *** | | no | no | *** | ** | * | no | *** | *** | ** |
| S71 | no | no | no | * | * | ** | *** | no | | no | ** | * | * | no | no | * | no |
| Y78 | * | *** | *** | ** | *** | *** | *** | no | no | | *** | * | no | no | *** | *** | ** |
| S87 | *** | *** | *** | no | * | no | ** | *** | ** | *** | | * | ** | *** | *** | *** | *** |
| T93 | *** | *** | *** | no | no | * | *** | ** | * | * | * | | no | ** | *** | *** | *** |
| L107 | *** | *** | *** | no | no | ** | *** | * | * | no | ** | no | | ** | *** | *** | *** |
| K124 | no | no | *** | *** | *** | *** | *** | no | no | no | *** | ** | ** | | ** | *** | no |
| E143 | *** | no | no | *** | *** | *** | *** | *** | no | *** | *** | *** | *** | ** | | *** | no |
| K157 | *** | *** | * | *** | *** | *** | *** | *** | * | *** | *** | *** | *** | *** | *** | | *** |
| G178 | no | no | *** | *** | *** | *** | *** | ** | no | ** | *** | *** | *** | no | no | *** | |

**Table S2.** The t-test results presented in **Figure 2** indicate a significant difference in the unbinding forces at different anchor points when pulled at a speed of 400 nm/s.

| Anchor points | $K_D$ dissociation constant [nM] |
| --- | --- |
| Q1 | 57.34 ± 14.84 |
| K62 | 37.85 ± 17.44 |
| K157 | 42.89 ± 12.92 |
| G178 | 42.52 ± 21.35 |

**Table S3.** Dissociation constants between CTLA-4 and anticalin at different anchor points.

| Anchor points | Δx [nm] (BE) | ln ($k_0$) [$s^{-1}$] (BE) | Δx [nm] (DHS) | ΔG [kJ/mol] (DHS) | ln ($\tau_0$) [s] (DHS) |
|---|---|---|---|---|---|
| Q1 | 0.37 ± 0.05 | -6.68 ± 1.54 | 0.54 ± 0.16 | 17.71 ± 1.62 | 7.42 ± 2.47 |
| Y32 | 0.46 ± 0.10 | -7.15 ± 2.50 | 0.38 ± 0.28 | 13.1 ± 4.03 | 3.25 ± 4.96 |
| I55 | 0.47 ± 0.02 | -12.69 ± 0.64 | 0.33 ± 0.15 | 18.53 ± 7.99 | 5.08 ± 2.30 |
| E57 | 0.78 ± 0.10 | -24.25 ± 3.97 | 0.31 ± 0.21 | 14.82 ± 3.37 | 4.62 ± 3.71 |
| K59 | 0.26 ± 0.02 | -5.27 ± 0.77 | 0.27 ± 0.08 | 18.74 ± 8.66 | 4.05 ± 1.18 |
| E60 | 0.33 ± 0.23 | -9.69 ± 9.91 | 0.34 ± 0.33 | 13.38 ± 5.95 | 4.06 ± 4.23 |
| K62 | 0.41 ± 0.13 | -15.42 ± 6.26 | 0.49 ± 0.48 | 15.75 ± 5.16 | 7.05 ± 7.31 |
| S63 | 0.58 ± 0.05 | -13.92 ± 1.55 | 0.39 ± 0.17 | 17.38 ± 4.28 | 5.1 ± 2.46 |
| S71 | 0.21 ± 0.05 | -0.89 ± 1.14 | 0.33 ± 0.22 | 15.51 ± 8.42 | 2.85 ± 2.94 |
| Y78 | 1.20 ± 1.05 | -28.34 ± 29.90 | 0.71 ± 0.31 | 17.5 ± 3.5 | 8.52 ± 4.03 |
| S87 | 0.39 ± 0.15 | -12.18 ± 6.59 | 0.36 ± 0.24 | 18.8 ± 4.74 | 7.09 ± 4.62 |
| T93 | 0.67 ± 0.04 | -19.44 ± 1.60 | 0.30 ± 0.13 | 19.72 ± 12.67 | 4.28 ± 1.99 |
| L107 | 0.34 ± 0.02 | -6.63 ± 0.82 | 0.49 ± 0.12 | 17.39 ± 2.36 | 7.64 ± 2.33 |
| K124 | 0.91 ± 0.69 | -19.78 ± 19.30 | 0.20 ± 0.06 | 8.92 ± 1.64 | 0.26 ± 1.42 |
| E143 | 0.55 ± 0.12 | -6.74 ± 2.56 | 0.25 ± 0.23 | 9.93 ± 3.66 | 0.92 ± 3.90 |
| K157 | 0.98 ± 0.13 | -11.99 ± 2.21 | 0.24 ± 0.19 | 8.97 ± 4.52 | 0.56 ± 3.62 |
| G178 | 0.32 ± 0.05 | -2.99 ± 1.14 | 0.64 ± 0.56 | 16.57 ± 5.05 | 6.59± 7.26 |

**Table S4.** Statistics of the energy barrier parameters including the distance to the energy barrier Δx (BE), the off rate at zero force $k_0$ (BE), Δx (DHS), the height of the energy barrier in the absence of force $\Delta G^{\ddagger}$ (DHS), and the zero-force life time $\tau_0$ (DHS) of Anticalin:(CTLA-4) complex at different pulling geometries.

| Anchor points | X position (Å) | Y position (Å) | Z position (Å) | Distance to CTLA4 mass center (Å) |
|---|---|---|---|---|
| CTLA-4 mass center | 60.29 | -17.66 | -23.67 | 0 |
| Q1 | - | - | - | - |
| Y32 | 84.19 | -27.96 | -10.33 | 29.25 |
| I55 | 79.58 | -34.97 | -8.45 | 30.06 |
| E57 | 79.06 | -40.99 | -6.98 | 34.28 |
| K59 | 77.82 | -47.39 | -7.54 | 38.10 |
| E60 | 78.28 | -51.05 | -6.64 | 41.58 |
| K62 | 81.00 | -49.78 | -11.18 | 40.21 |
| S63 | 78.92 | -47.15 | -12.95 | 36.49 |
| S71 | 72.63 | -22.20 | -9.46 | 19.36 |
| Y78 | 70.52 | -30.98 | -9.52 | 21.96 |
| S87 | 85.17 | -51.20 | -22.96 | 41.76 |
| T93 | 78.28 | -44.48 | -22.02 | 32.34 |
| L107 | 81.02 | -41.72 | -25.25 | 31.80 |
| K124 | 80.78 | -37.19 | -27.68 | 28.59 |
| E143 | 90.62 | -22.56 | -9.49 | 33.84 |
| K157 | 92.58 | -26.39 | -30.46 | 34.13 |
| G178 | 76.60 | -27.64 | -1.11 | 29.57 |

**Table S5.** Distance between different anticalin anchor points and CTLA-4 mass center calculated from Anticalin:(CTLA-4) complex structure in PDB file 3BX7.

| | Y | $X_1$ | $X_2$ | $X_3$ | $X_4$ | |
|---|---|---|---|---|---|---|
| Anchor points | Unbinding force @400 nm/s [pN] | Hydropathy | Electronic charge index | Solvent accessible surface area | Distance to CTLA-4 mass center | Secondary structure |
| Q1 | 128.8 | −0.69 | 1.36 | - | - | N/C terminal |
| Y32 | 107.2 | 0.02 | 0.72 | 22.42 | 29.25 | β strand |
| I55 | 160.5 | 0.73 | 0.09 | 29.26 | 30.06 | β strand |
| E57 | 163.1 | −0.62 | 1.31 | 71.06 | 34.28 | β strand |
| K59 | 161.6 | −1.10 | 0.53 | 90.81 | 38.10 | Loop |
| E60 | 205.7 | −0.62 | 1.31 | 186.77 | 41.58 | Loop |
| K62 | 218.9 | −1.10 | 0.53 | 133.37 | 40.21 | Loop |
| S63 | 138.7 | −0.26 | 0.56 | 2.07 | 36.49 | Loop |
| S71 | 98.7 | −0.26 | 0.56 | 26.12 | 19.36 | β strand |
| Y78 | 122.4 | 0.02 | 0.72 | 122.66 | 21.96 | β strand |
| S87 | 176.1 | −0.26 | 0.56 | 117.06 | 41.76 | Loop |
| T93 | 154 | −0.18 | 0.65 | 38.57 | 32.34 | β strand |
| L107 | 146.1 | 0.53 | 0.10 | 36.21 | 31.80 | β strand |
| K124 | 112.1 | −1.10 | 0.53 | 38.80 | 28.59 | β strand |
| E143 | 91.8 | −0.62 | 1.31 | 129.20 | 33.84 | Loop |
| K157 | 72.4 | −1.10 | 0.53 | 103.17 | 34.13 | α helix |
| G178 | 107.8 | 0.16 | 0.02 | 81.54 | 29.57 | N/C terminal |

**Table S6.** Statistics of the unbinding force at a pulling speed of 400nm/s, and the relative position, hydropathy, electronic charge index, solvent accessible surface area, and distance to CTLA4 mass center of the original residue where we introduced anchor points at different pulling geometries.

| | Y (B) | $X_1$ (D) | $X_2$ (E) | $X_3$ (F) | $X_4$ (G) | |
|---|---|---|---|---|---|---|
| Anchor points | Unbinding force @400 nm/s [pN] | Hydropathy | Electronic charge index | Solvent accessible surface area | Distance to CTLA4 mass center | Secondary structure |
| Y32 | 107.2 | 0.02 | 0.55 | 0.12 | 0.70 | β strand |
| I55 | 160.5 | 0.66 | 0.07 | 0.16 | 0.72 | β strand |
| E57 | 163.1 | -0.56 | 1.00 | 0.38 | 0.82 | β strand |
| K59 | 161.6 | -1.00 | 0.40 | 0.49 | 0.91 | Loop |
| E60 | 205.7 | -0.56 | 1.00 | 1.00 | 1.00 | Loop |
| K62 | 218.9 | -1.00 | 0.40 | 0.71 | 0.96 | Loop |
| S63 | 138.7 | -0.24 | 0.43 | 0.01 | 0.87 | Loop |
| S71 | 98.7 | -0.24 | 0.43 | 0.14 | 0.46 | β strand |
| Y78 | 122.4 | 0.02 | 0.55 | 0.66 | 0.53 | β strand |
| S87 | 176.1 | -0.24 | 0.43 | 0.63 | 1.00 | Loop |
| T93 | 154 | -0.16 | 0.50 | 0.21 | 0.77 | β strand |
| L107 | 146.1 | 0.48 | 0.08 | 0.19 | 0.76 | β strand |
| K124 | 112.1 | -1.00 | 0.40 | 0.21 | 0.68 | β strand |
| E143 | 91.8 | -0.56 | 1.00 | 0.69 | 0.81 | Loop |
| K157 | 72.4 | -1.00 | 0.40 | 0.55 | 0.82 | α helix |
| G178 | 107.8 | 0.15 | 0.02 | 0.44 | 0.71 | N/C terminal |

**Table S7.** Normalized parameters from Table S6 for the multi-parameter linear regression fitting. Anticalin with anchor point Q1 was omitted due to insufficient data in the solvent accessible surface area and the distance to CTLA-4 mass center. Each feature was normalized by the absolute maximum number in its group as described in Methods.
